# Characterization of evolutionarily conserved key players affecting eukaryotic flagellar motility and fertility using a moss model

**DOI:** 10.1101/728691

**Authors:** Rabea Meyberg, Pierre-François Perroud, Fabian B. Haas, Lucas Schneider, Thomas Heimerl, Karen Renzaglia, Stefan A. Rensing

## Abstract

Defects in flagella/cilia are often associated with infertility and disease. Motile male gametes (sperm cells) with flagella are an ancestral eukaryotic trait that has been lost in several lineages, for example in flowering plants. Here, we made use of a phenotypic male fertility difference between two moss (*Physcomitrella patens*) strains to explore spermatozoid function. We compare genetic and epigenetic variation as well as expression profiles between the Gransden and Reute strain to identify a set of genes associated with moss male infertility. Defects in mammal and algal homologs of these genes coincide with a loss of fertility, demonstrating the evolutionary conservation of flagellar function related to male fertility across kingdoms. As a proof of principle, we generated a loss-of-function mutant of a coiled-coil domain containing 39 (ccdc39) gene that is part of the flagellar hydin network. Indeed, the *Ppccdc39* mutant resembles the male infertile Gransden strain phenotype. Potentially, several somatic (epi-)mutations occurred during prolonged vegetative propagation of *P. patens* Gransden, causing regulatory differences of e.g. the homeodomain transcription factor BELL1. Probably these somatic changes are causative for the observed male fertility. We propose that *P. patens* spermatozoids might be employed as an easily accessible system to study male infertility of human and animals.

## Introduction

### Motile plant gametes allow studying sperm cell fertility

Motile (flagellated) gametes are an ancestral character of eukaryotes (Mitchell 2007, Stewart *et al.* 1975) that has been secondarily lost in lineages such as the Zygnematales (Transeau 1951), which reproduce via conjugation of aplanogametes, and flowering plants, in which pollen transport non-motile gametes to the female (Renzaglia *et al.* 2001, Southworth *et al.* 1997). Flagella play an important role in sexual reproduction. In the unicellular algal model organism *Chlamydomonas reinhardtii* flagella are responsible for motility of the organism and serve during sexual reproduction to mediate a species-specific adhesion between two cells of different mating types (van den Ende *et al.* 1990). In streptophyte algae (Streptophyta comprise charophycean green algae as well as land plants), male gametes (spermatozoids) are the only motile cells of sessile multicellular algae, swimming to the oogonia to fertilize the egg cell (Hackenberg *et al.* 2019, McCourt *et al.* 2004). After the water-to-land-transition of plants this system was retained, i.e. motile (flagellated) spermatozoids require water to swim to the egg cell. Flagella of spermatozoids from flagellated organisms show a common architecture (Carvalho-Santos *et al.* 2011) and were already present in the most recent common ancestor (MRCA) of land plants (Stewart *et al.* 1975). During land plant evolution, loss of motile sperm occurred in seed plants, after the evolution of cycads and *Ginkgo* (both of which have pollen and motile male gametes), probably in the MRCA of conifers, Gnetales and flowering plants (Renzaglia *et al.* 2000, Renzaglia *et al.* 2001). Flagellated plants, that have retained motile spermatozoids, are an easily accessible system to study the fertility and other characteristics of male gametes. The moss *Physcomitrella patens* is particularly attractive in that regard because it develops spermatozoids in superficial male sex organs (antheridia) on an independent generation (gametophyte) that is readily cultured and manipulated (Cove 2005, Landberg *et al.* 2013).

### The moss model Physcomitrella patens

Bryophytes comprise the mosses, liverworts and hornworts and probably represent the monophyletic sister clade to vascular plants (Puttick *et al.* 2018). Due to this informative phylogenetic position they are increasingly being used to address evolutionary-developmental questions, e.g. (Aya *et al.* 2011, Horst *et al.* 2016, Sakakibara *et al.* 2013, Sakakibara *et al.* 2008), but also for functional studies of mammalian homologs in easily accessible model organisms (Ortiz-Ramirez *et al.* 2017, Sanchez-Vera *et al.* 2017). *P. patens* is a well-developed functional genomics moss model system. The genome was published in 2008 (Rensing *et al.* 2008) and recently updated to chromosome scale (Lang *et al.* 2018). Collections of worldwide accessions are available (Beike *et al.* 2014), of which some are likely to constitute ecotypes (Lang *et al.* 2018). The predominantly used ecotype Gransden (Gd) was derived from a single spore isolate collected 1962 in Gransden Wood (UK). The Gd cultures used in most labs worldwide were typically propagated vegetatively and several labs reported fertility issues over the past decades (Ashton *et al.* 2000, Hiss *et al.* 2017, Landberg *et al.* 2013, Perroud *et al.* 2011), potentially the result of somatic (epi-)mutations through decades of vegetative propagation (Ashton *et al.* 2000). The ecotype Reute (Re) collected 2006 in Reute (Germany) has a low genetic divergence to Gd, is highly self-fertile and thus has recently been introduced as a Gd alternative to enable studies involving sexual reproduction (Hiss *et al.* 2017).

### P. patens sexual reproduction

*P. patens’* sexual reproduction is initiated (under natural conditions in autumn) when day length shifts towards short days and temperature drops (Engel 1968, Hohe *et al.* 2002, Nakosteen *et al.* 1978). On the tip (apex) of the leafy gametophores the apical stem cell produces the antheridium initial stem cell (Kofuji *et al.* 2018), which subsequently gives rise to a bundle of antheridia, the male reproductive organs (Fig. 1). Each antheridium consists of a single outer cell layer or jacket surrounding a mass of spermatogenous tissue that produces up to 160 motile spermatozoids. Upon maturity the tip cell of the antheridia swells and bursts (when moistened by water) to release mature, swimming male gametes (spermatozoids). A few days after initiation of antheridia development, the female gametangia (archegonia) start to develop (Kofuji *et al.* 2009). They comprise a flask shaped egg-containing venter and elongated neck that opens following degeneration of the neck canal cells upon maturity, allowing swimming spermatozoids to reach the egg cell (Hiss *et al.* 2017, Landberg *et al.* 2013, Sanchez-Vera *et al.* 2017). After fertilization, the diploid zygote undergoes embryogenesis and develops into the sporophyte, which eventually generates haploid spores via meiosis (Fig. 1). Considering the difference in fecundity of Gd and Re (Hiss *et al.* 2017) we aim here to determine the genes underlying this difference, employing *P. patens* as a model for studying sperm fertility. For this purpose, we analysed the nearly self-sterile ecotype Gransden (Gd) in comparison with the highly fertile ecotype Reute (Re), using genetic and epigenetic variation data, fertility phenotyping as well as transcript profiling. Based on publicly available array data of developmental stages during sexual reproduction and network analyses, we generated a deletion mutant of a gene exclusively expressed in sexual reproductive organs. The mutant displays a loss of male fertility similar to that observed in Gd, and to that in human/mammal mutants of the ortholog. Our study demonstrates that the moss male germ line can serve as a model for flagellar dysfunction disease.

**Figure 1:**
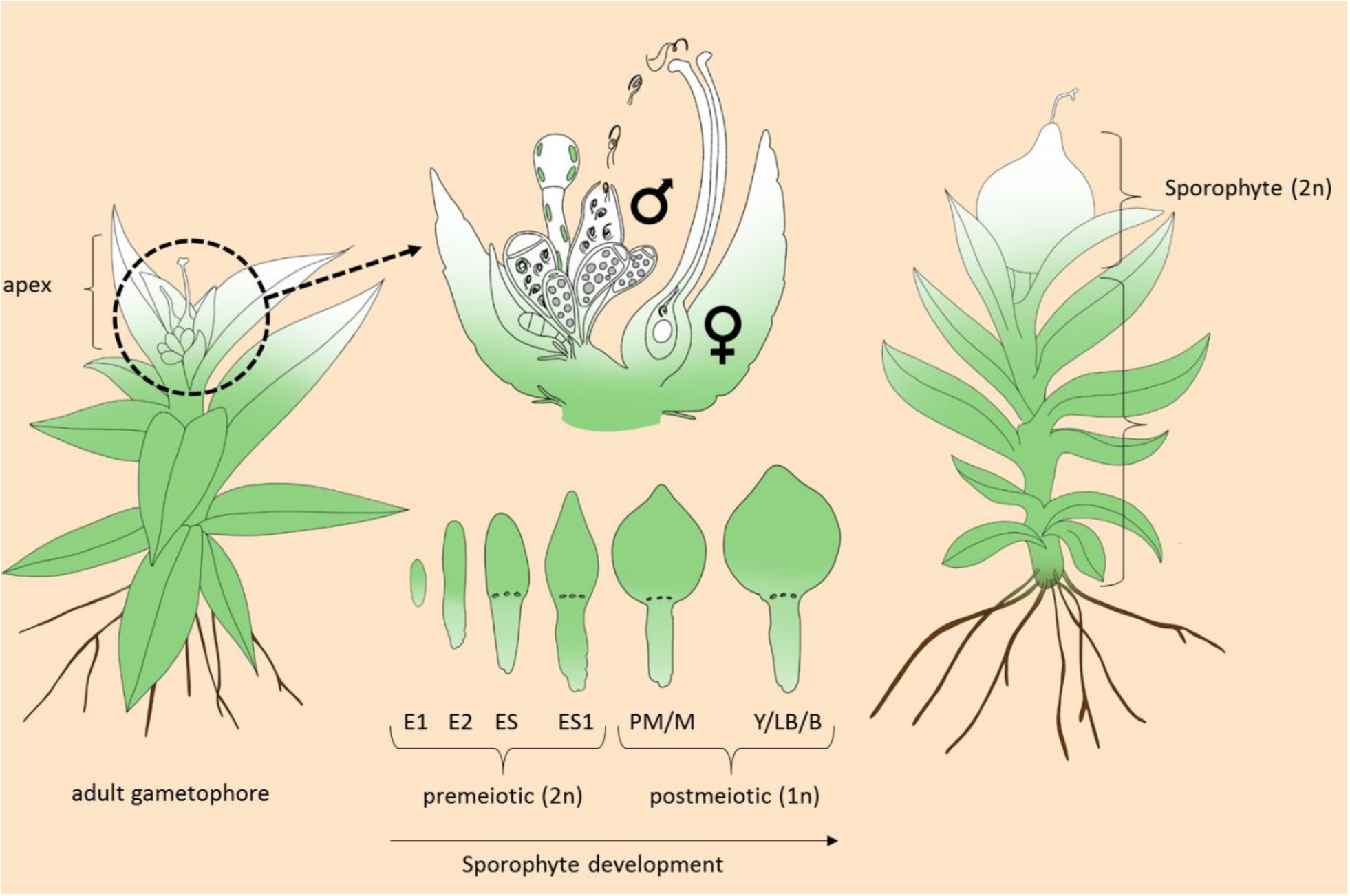
Illustration of *P. patens* sexual reproduction. Upon environmental stimulus, reproductive organs (gametangia) develop on the apex of each gametophore, archegonia (female) and antheridia (male). Mature antheridia release the motile (flagellated) spermatozoids upon watering. The sperm cells swim through the archegonial venter to fertilize the egg cell. After fertilization, embryo development (E1/E2) and sporophyte development (ES-B) occurs. The mature sporophyte is located on the apex of the gametophore and releases haploid spores of the next generation. *P. patens* is predominantly selfing. Embryo/sporophyte developmental stages according to Hiss et al. (2017).

## Results & Discussion

### The Gransden ecotype displays a reduced male fertility and a defect in spermatozoid motility

The ecotype Reute (Re) was shown to develop a significantly higher number of sporophytes per gametophore in comparison to the broadly used ecotype Gransden (Gd, (Hiss *et al.* 2017)). To determine whether the reduced fertility of Gd is based on the female or male reproductive apparatus, crossing analyses between Gd and Re were performed, using the fluorescent marker strain Re-mCherry (Perroud *et al.* 2011, Perroud *et al.* 2019). Re developed on average 100% sporophytes per gametophore, of which 0.79% were crosses (Fig. 2A).

**Figure 2:**
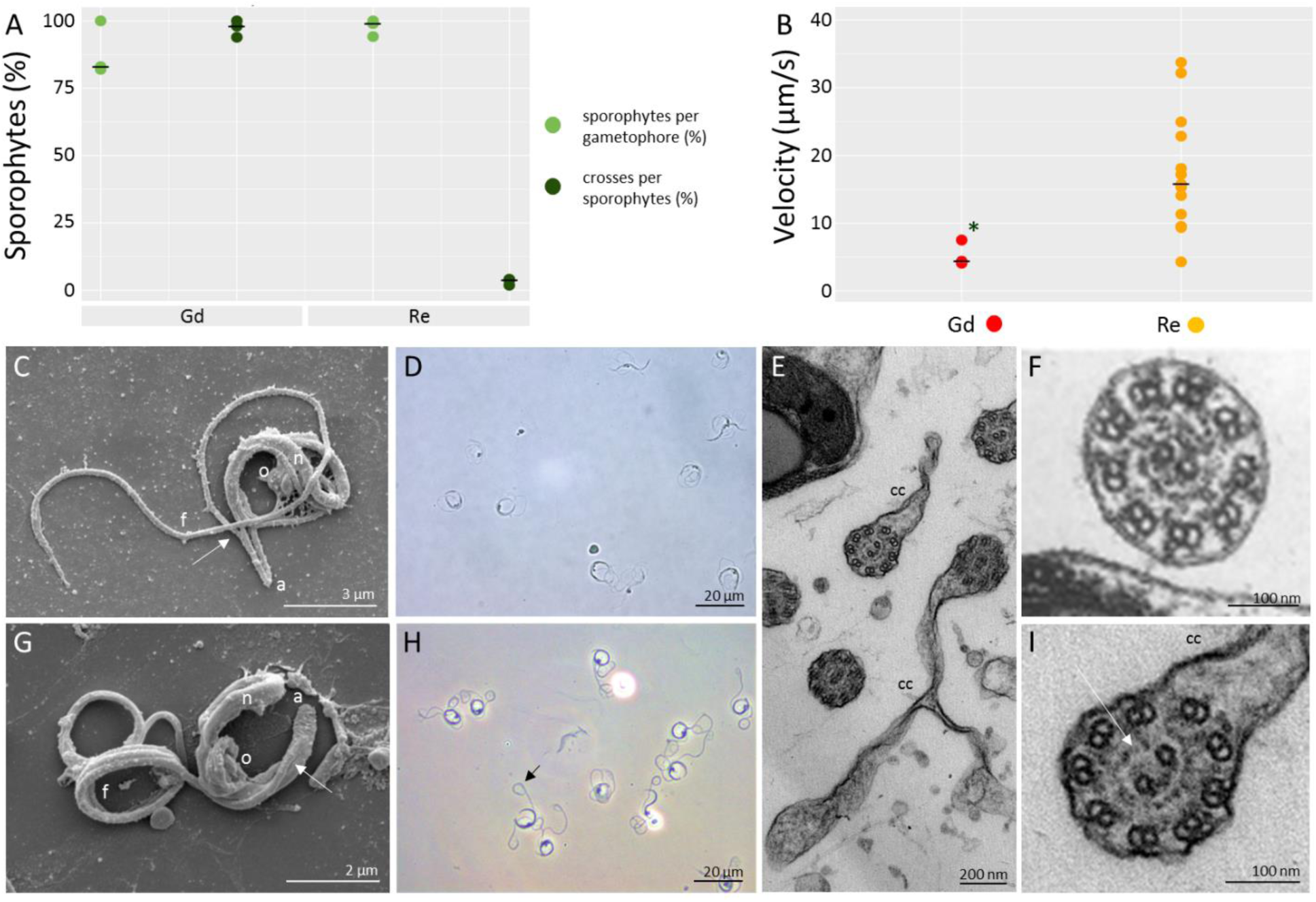
Comparative analysis of Gd and Re male reproductive apparatus. A: Crossing analysis between Re (n = 430), Gd (n = 300) and fluorescent marker strain Re-mcherry. Re develops 100% of sporophytes per gametophore of which 0.79% are crosses. Gd develops 88.13% sporophytes per gametophore of which 99.69% are crosses. Median represented by black line. Significance shown by asterisk (Chi-square test, p < 0.01). B: Re (n = 10) and Gd (n = 3) spermatozoids differ significantly in velocity (two-sided t-test, p < 0.01,*). Median represented by black line. C: SEM of Re sperm cell showing cell architecture. From the cell anterior (a), the two flagella emerge (arrow) at staggered locations from the coiled nucleus (n). Organelles (o) include one mitochondrion and one plastid that attach to the mid-section of the nucleus. One of two long flagella (f) is visible and extends beyond the cell proper. D: Phase contrast image of swimming Re spermatozoids. E: TEM of mature Gd axonemes. Cytoplasmic connections (cc) appear on irregular flagella. F: TEM of a mature spermatozoid of Re. The axoneme exhibits clear outer dynein arms and the plasmalemma is closely associated with the nine doublets. G: SEM of Gd spermatozoid with architecture as in Re spermatozoids in C, except the flagella remain coiled. H: Phase contrast image of swimming Gd spermatozoids showing loops at the posterior end of the flagella (arrow). I: TEM of mature axonemes of Gd spermatid showing missing parts of the central pair microtubuli projections (white arrow) and cytoplasmic connections (cc).

This rate is expected, since *P. patens* is known to predominantly self-fertilize (Perroud *et al.* 2011). In comparison, Gd developed 88.13% sporophytes per gametophore of which 99.69% were crosses (significantly more than Re: p < 0.01, t-test, Fig. 2A). Thus, Gd archegonia are fully functional and can develop comparable numbers of sporophytes when fertilized by a male fertile partner. Hence, the Gd male reproductive apparatus is impaired. Analysis of the spermatozoid number per antheridium showed no significant differences between Re (median 123) and Gd (median 125, Figure S1A). This is less than determined by an approximation method (Horst *et al.* 2017), counting DAPI stained spermatozoids not released from the antheridium. Motility measurements showed that only a small number of Gd spermatozoids are motile (Fig. S1B), and show a significantly reduced velocity (Re median: 15,75 µm/s; Gd median: 4,37µm/s; p < 0.01, t-test; Fig. 2B). After release of the spermatozoids through the bursting tip cells of the antheridia, Re spermatozoids started moving a few seconds after release, whereas Gd spermatozoids showed motility only rarely. With regard to their structure, Gd spermatozoids display significantly more spermatozoids with the ends of flagella remaining in a coil (90.2%) than Re (6.6%, t-test, p < 0,01, Fig. 2C, D, G, H, Fig. S1C). Interestingly, similar coiled flagella are also known from mouse and human infertile flagella (Dong *et al.* 2018, He *et al.* 2018, Tang *et al.* 2017). There are ultrastructural differences between Re and Gd sperm cells. Two cylindrical flagella develop around the outside of the spermatozoid as the nucleus condenses and elongates (Fig. 2C, D, G, H). The axonemes of both ecotypes exhibit the typical nine doublets and two central pair microtubules that characterize flagella of most eukaryotes (Fig. 2E, F, I). Axonemes in Re mature spermatozoids demonstrate visible outer dynein arms and protein projections at the central pair of microtubuli (Fig. 2F). In contrast to Re (Fig. 2C, D), gametes of Gd appear to arrest in the final stages of flagellar elongation and thus the posterior coils of the flagella fail to individualize, resulting in the coiled posterior loop (Fig. 2G, H). Mature Gd axonemes developed cytoplasmic connections, which probably result in the coiled posterior loop of the flagella. Additionally the protein projections around the central pair of microtubules seems to lack some proteins (Fig. 2 E,I).

### Genes related to flagellar assembly and motility harbour (epi-)mutations between Gd and Re

Similar to motile sperm, DNA methylation is an ancestral eukaryotic feature (Feng *et al.* 2010) and is supposed to regulate gene expression on the DNA level (Zemach *et al.* 2010). In *P. patens*, methylated gene bodies usually are associated with lower gene expression (Lang *et al.* 2018) and loss of the DNA methyltransferase PpMET1, which is involved in CG DNA methylation of gene bodies, inhibits sporophyte development (Yaari *et al.* 2015). Recently, it has been shown, that the male reproductive organs undergo severe changes in DNA methylation in the liverwort *M. polymorpha* during sexual reproduction (Schmid *et al.* 2018). Hence, we expected DNA methylation to play a role during regulation of sexual reproduction in moss. Whole genome bisulfite sequencing (bs-seq) of Re and Gd adult gametophores (bearing gametangia; Fig. 1) was performed. Differentially methylated positions (DMPs) in all three methylation contexts (CHG, CHH, CG) were determined. In total, 671 genes harboring DMPs were found (Table S1). GO bias analysis of the genes containing DMPs showed enriched terms related to cilium movement and motile cilium assembly, as well as protein and macromolecule modification, which includes post-translational modifications like ubiquitination (Fig. S2A). Using the intersect of genes that contain DMPs and single nucleotide polymorphisms (SNPs, Table S2, (Hiss *et al.* 2017)) the number of GO terms associated with cilia and microtubule based movement was found to be increased (from 3 to 6, Fig. 3A), suggesting genetic and epigenetic effects on the Gd phenotype. In the intersection, additional GO terms related to protein phosphorylation were found to be over-represented (Fig. 3A). In *C. reinhardtii*, it was shown that protein phosphorylation is a key event of flagellar disassembly (Pan *et al.* 2011) and that the phosphorylation state of an aurora-like protein kinase coincides with reduced flagellar length (Luo *et al.* 2011).

**Figure 3:**
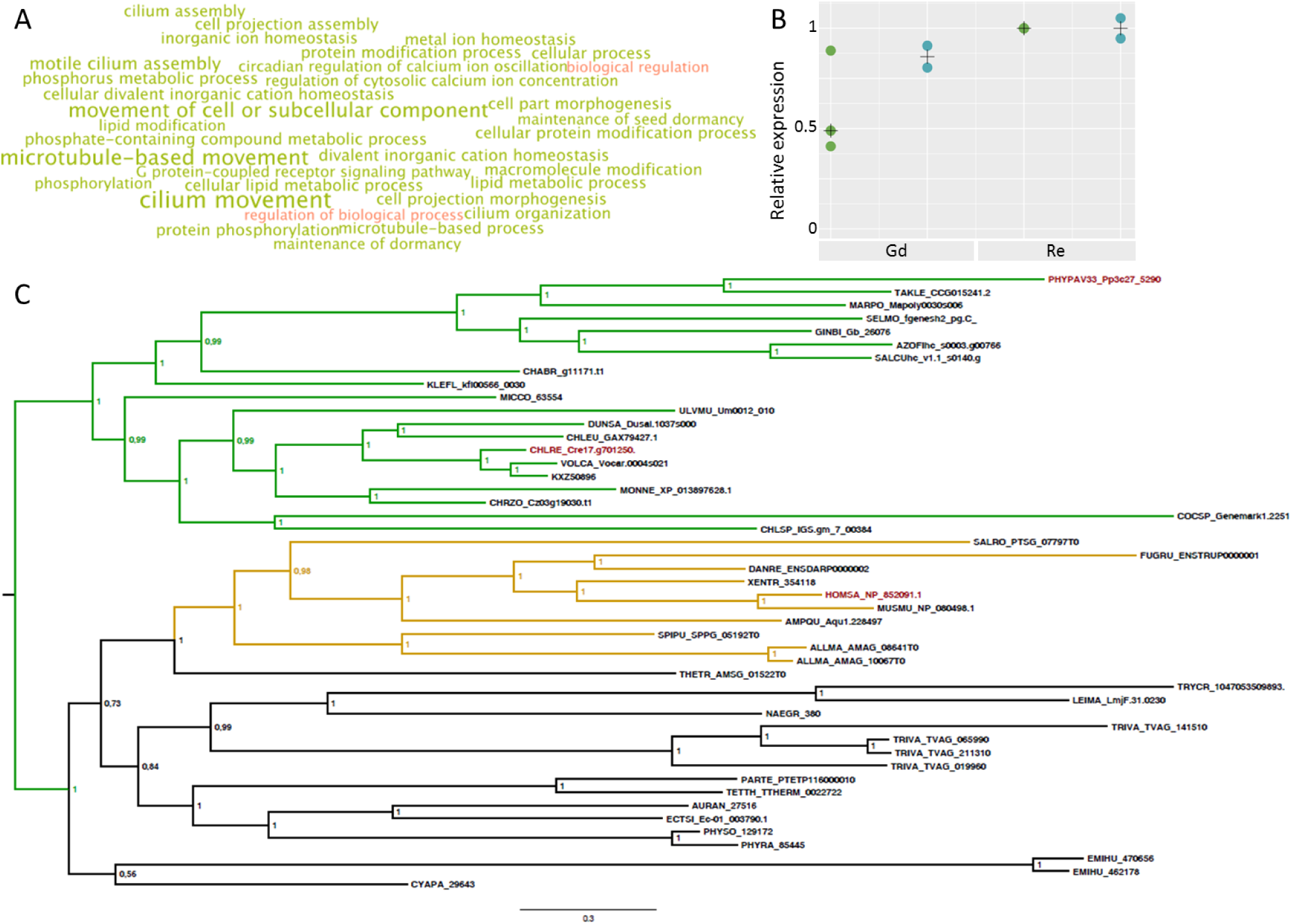
DMP/SNP overlap and CCDC39. A: GO bias analysis of genes differentially methylated and SNP-containing in comparison of Gd vs. Re. The intersect of DMP and SNP affected genes shows over-representation of GO terms associated with cilia motility. Over-represented terms are shown in green, whereas under-represented terms are shown in red. Larger font size correlates with a higher significance level. B: Relative expression of ccdc39 in qPCR (green, n = 3) and RNA-seq (blue, n = 2) data shows higher expression in Re in comparison to Gd. Median marked by black cross. C: Phylogenetic analysis of CCDC39 shows clear correlation with the presence of flagella and generally reflects species relationships. Planta are shown in green, opisthokonts in yellow, protozoa, SAR and fungi in black. *Homo sapiens, Chlamydomonas reinhardtii* and *Physcomitrella patens* are marked in red.

### RNA-seq of male reproductive organs suggests differential transcriptional regulation to be connected to infertility

To gain deeper insight into gene expression differences, cDNA sequencing (RNA-seq) of Gd and Re antheridial bundles was performed. In total 19,218 genes were found to be expressed (RPKM >= 2) in Gd and 19,254 in Re (Table S3/S4). 865 genes were uniquely expressed in Gd and 901 genes in Re (Table S5/S6). GO bias analysis of all genes expressed in antheridia of Gd (Fig. S2B) and Re (Fig. S2C) showed an over-representation of terms like amide biosynthetic process and peptide biosynthetic process. Spermine and spermidine belong to the amides and play a major role in male fertility in mammals (Lefevre *et al.* 2011) which, based on the RNA-seq GO analysis, also appears to be the case in *P. patens*. The few under-represented terms found are all connected to mRNA capping, potentially indicating that during gamete development mRNA capping could be performed by less characterized capping enzymes, or that translation occurs independently of mRNA capping. It was postulated that the first eukaryotic mRNA translation was probably driven by internal ribosomal entry sites (IRES) and that the cap-dependent (CD) mechanism evolved later. The former mechanism, according to this hypothesis, was retained as an additional regulation of the translation in specific situations like stress responses (Godet *et al.* 2019, Hernandez 2008). Also, cap-independent (CI) translation was recently suggested to be a robust partner of CD translation and to appear prevalently in germ cells (Keiper 2019), which matches with the presented results.

In human, transcriptional regulation is important to control spermatogenesis (Bettegowda *et al.* 2010). Within the expressed genes, in total 1,259 transcription associated proteins (TAPs; comprising transcription factors, TFs, and transcriptional regulators, TRs) could be identified using TAPscan (Wilhelmsson *et al.* 2017). Within the Re uniquely expressed genes, in total 74 TAPs of 27 different TAP families could be identified, including HD_BEL and MADS_MIKC^c^ type proteins (Table S7). Interestingly, bell1 (HD_BEL) previously was described not to be expressed in antheridia (Horst *et al.* 2016), which indeed is true for Gd (used in the study by Horst *et al.*), but not for Re (Fig. S3A). Therefore, BELL1 might play a role in male fertility in *P. patens*. MADS-box genes are important for flower, pollen and fruit development (Theißen *et al.* 2016) and have previously been associated with sexual reproduction in *P. patens* (Hohe *et al.* 2002, Quodt *et al.* 2007, Singer *et al.* 2007). *P. patens* MIKC^c^-type genes are important for motile flagella and external water conduction (Koshimizu *et al.* 2018). In the latter study, PPM1, PPM2 and PPMC6 are the key players for flagellar motility, but since the triple KO did not completely phenocopy the sextuple KO (*ppm1, ppm2, ppmads1, ppmc5, ppmc6, ppmadss*), the other genes apparently also play a role. While ppm1, ppm2 and ppmc6 are expressed in both, Re and Gd antheridia, mads1 (Hohe *et al.* 2002) and ppmc5 are exclusively expressed in Re, implying a putative involvement of these two genes in flagellar motility. Among the Gd uniquely expressed genes (Table S5), 40 TAPs belonging to 23 different TAP families were found, including MADS and HD proteins. TIM1 (Pp3c17_24040V3.1) is a type I MADS-box protein, basal to all other type I MADS-box proteins (Gramzow *et al.* 2010), whereas the HD gene duxa-like (Pp3c5_6470V3.1) is mainly expressed in lung and testis tissue in human, which supports a potential function in cilia/flagella (Booth *et al.* 2007).

### Analysis of differentially expressed genes between Gd and Re

From all expressed genes in Gd and Re, a set of 110 differentially expressed genes (DEGs) could be identified using a stringent intersect of three tools. Detailed analysis of the DEGs showed many spermatozoid/sperm and pollen related genes (Table S8). Of the genes expressed significantly higher in Re, many showed no expression in Gd, but moderate to high expression in Re, e.g. the arl13b homolog (Pp3c1_40600V3.1), which plays a role in flagella and cilia stability and signaling via axonemal poly-glutamylation, and the protein phosphatase hydin homolog Pp3c3_14230V3.1 (Fig. S3B). The central microtubule pair protein hydin is a well analysed gene in human and algae, and connected to the human disease Primary Ciliary Dyskinesia (PCD). Mutations in this gene lead to the loss of flagellar motility and occasionally to loss of one or both central pair microtubules due to missing C2b projections (Lechtreck *et al.* 2007, Olbrich *et al.* 2012). Interestingly, the Gd ultrastructure showed missing protein projections (Fig. 2E,I) which could might result from the missing hydin expression.

Genes that are expressed significantly higher in Gd often lacked annotation (28/37 showed no annotation in the latest gene annotation v3.3 (Lang *et al.* 2018)) and hence might be lineage specific. Among the annotated genes more highly expressed in Gd were e.g. the membrane associated ring finger march1 homolog (Pp3c18_16700V3.1), which is known to negatively affect the sperm quality and quantity in Chinese Holstein bulls (Liu *et al.* 2017), and an arabinogalactan 31 homolog (Pp3c5_9210V3.1). Arabinogalactan proteins are known to be part of mucilage (Lord *et al.* 1992). Analysis of the mucilage content during antheridia maturation in the charophytic alga *Chara vulgaris* showed a reduction of mucilage upon maturity (Gosek *et al.* 1991). Arabinogalactan proteins are abundant in the matrix around fern spermatozoids during development and are virtually negligible when antheridia release spermatozoids (Lopez *et al.* 2018). Thus, an overexpression of an arabinogalactan gene could represent an impairment of the spermatozoid maturation process in the Gd background.

RNA-seq expression data of hydin and march1 have been confirmed by RT-PCR (Fig. S3B), and SNP and DNA methylation data have been analysed. The hydin gene body and potential promoter region (2kbp upstream of the coding sequence) displays six SNPs and one insertion or deletion (InDel) between Gd and Re. One SNP is located in the putative promoter region, three are intron variants and two are located within an exon, causing moderate amino acid changes (Table S9). In Gd, CHG and CHH context DMPs are present in the promoter and gene body, while a single CG mark is present in Re (Fig. S4). The lack of expression in Gd matches the presence of gene body methylation, shown to coincide with lack of expression (Lang *et al.* 2018). March1 does show very low levels of DNA methylation but two SNPs between Gd and Re in the 5’-UTR and five SNPs and one InDel in the putative promoter region, which could affect a potential regulatory function of the UTR (Fig. S5, Table S9). Network analysis was performed to gain more insights into putative protein interaction partners. The March1 network revealed proteins known to be involved in degradation of mis-folded proteins via ubiquitination, and modification of proteins via phosphorylation (Fig. S6A). Since protein phosphorylation and ubiquitination pathways are known to act together (Hunter 2007), march1 disregulation in Gd might be involved in the determined phenotype. The hydin network revealed many proteins involved in ciliary function and motility e.g. radial spoke head protein 9 (RSPH9, (Castleman *et al.* 2009) and CCDC39 (Merveille *et al.* 2011, Oda *et al.* 2014), Fig. S6B.

CCDC39 (Pp3c27_5290V3.1) was detected via a candidate gene approach because it shows expression only in adult gametophores (bearing gametangia), and a difference between Re and Gd (Fig. 3B). It is part of the hydin network, is a coiled-coil domain containing protein which, as well as hydin and rsph9, belongs to the genes that cause PCD when mutated (Antony *et al.* 2013, Horani *et al.* 2018). In human it is required for the assembly of inner dynein arms and the dynein regulatory complex (Merveille *et al.* 2011) and in *Chlamydomonas reinhardtii* it acts as a molecular ruler for the determination of the flagellar length (Oda *et al.* 2014). Phylogenetic analysis showed the presence of ccdc39 orthologs in all major kingdoms, providing evidence that it was already present in the MRCA of all eukaryotes (Fig. 3C). Interestingly, all species with ccdc39 orthologs also possess flagella, implicating a gene function unique to these motile organelles. Ccdc39 does not show amino acid sequence-affecting SNPs between Gd and Re in the gene body, and no SNPs in the promoter region (Table S9). The promoter region and gene body of ccdc39 display low levels of DNA methylation (Fig. S6). Interestingly, CCDC39 and MARCH1 network analysis showed connections to the same two protein phosphatases (Pp3c16_18360V3.1, Pp3c25_7050V3.1, Fig. S6A, C), implicating involvement of phosphatases in spermatogenesis of *P. patens*.

### Ccdc39 is essential for proper flagellar development in P. patens

Expression analysis of ccdc39 via RNA-seq and qPCR in *P. patens* showed no expression in any tissue except for the antheridia, and a reduced expression level in Gd as compared to Re (Fig. 3C), suggesting a functional connection to the observed fertility phenotype. To elucidate the function of ccdc39 and to separate the phenotype from hydin function, a loss-of-function mutant was generated, hereafter referred to as *ccdc39*. Protoplast regeneration, protonemal and gametophytic growth and gametangia development did not reveal an obvious aberrant phenotype (Fig. S8, S9). However, no sporophytes were formed under selfing conditions (Fig. 4A, B yellow). 30 days after watering Re had developed mature brown sporophytes, whereas *ccdc39* apices showed several unfertilized archegonia and bundles of antheridia in a cauliflower-like structure (Fig. 4C). When no fertilization takes place, *P. patens* is known to continue gametangiogenesis, leading to the accumulation of multiple archegonia per gametophore (Landberg *et al.* 2013, Sanchez-Vera *et al.* 2017). Under crossing conditions using the male fertile Re-mCherry line (Perroud *et al.* 2019) a close to normal amount of sporophytes (94%) could be observed, with 100% of them being crosses (Fig. 4A, light and dark green), suggesting a defect in the male reproductive apparatus. Variable phenotypes of flagella could be detected in *ccdc39*. Some spermatozoids developed very short flagella, others normal length flagella, but with ends that remained in coils, comparable to the Gd phenotype (Fig. 4D-F). Pp*ccdc39* hence showed a somewhat different phenotype compared to the alga *C. reinhardtii*, in which a severe reduction of the flagellar length occurred (Oda *et al.* 2014), an observation only occasionally seen in *P. patens ccdc39*. No *ccdc39* null mutant has been reported for *H. sapiens* yet, but cilia of epithelial cells of persons bearing mutations in ccdc39 showed a reduction of length in 2/4 cases (Merveille *et al.* 2011), implying flagellar length variations to be part of the phenotype, comparative to *P. patens* ccdc39. Interestingly, all three species share the loss of the inner arm dyneins upon mutation of ccdc39 ((Merveille *et al.* 2011, Oda *et al.* 2014); Fig.4 G-I), indicating a conserved function. Ccdc39 gametes undergo normal cellular morphogenesis into a streamlined coiled architecture, including nuclear compaction/elongation, similar to Re and Gd gametes (Fig. 4F). The locomotory apparatus develops normally and the two flagella elongate around the cell perimeter. However, axonemal organization is completely disrupted in the KO spermatozoids. The nine doublet microtubules are randomly arranged within an abnormally-shaped flagellar shaft that is rarely cylindrical (Fig. 4E-I). The central pair complex remains intact but it is not anchored in the center of the axoneme (Fig. 4F-I). Floating doublet microtubules show evidence of outer dynein but inner dynein is unclear (Fig. 4G, H). Typically, only a single complement of nine doublets and one central pair occurs within an axoneme, but the occasional two complements are visible within a single flagellar shaft (Fig. 4F, I). Cytoplasmic connections across coils provide evidence of incomplete separation of cellular coils that is likely responsible for the posterior loop in flagella evident in the light microscope (Fig. 4D). The cytoplasmatic connections observed in the *ccdc39* background are similar to those displayed by Gd (Fig. 2E), which also showed a reduced expression of ccdc39 in comparison to Re (Fig. S3B). Together these observations suggest a functional connection between ccdc39 expression and the mutant phenotype.

**Figure 4:**
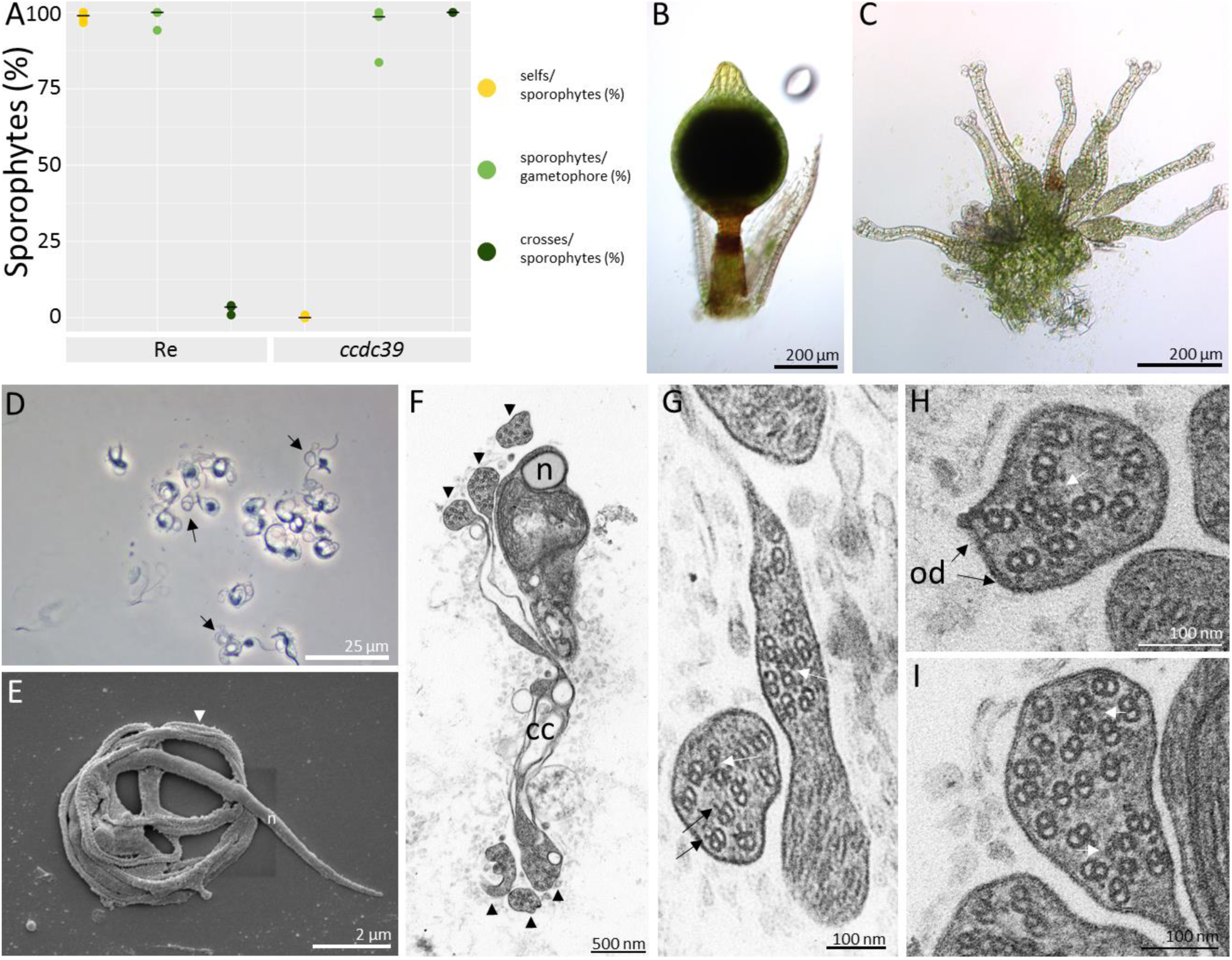
Analysis of *P. patens ccdc39*. A: Selfing and crossing analysis between *ccdc39* (n = 630), Re (n = 331) and fluorescent marker strain Re-mcherry. *ccdc39* develops 0% and Re develops 99% sporophytes per gametophore under selfing conditions. Under crossing conditions, Re develops 98% of sporophytes per gametophore of which 0.03% are crosses. *ccdc39* develops 94% sporophytes per gametophore of which 100% are crosses (asterisk shows significant deviation, Chi-square test, p < 0.01). Median marked by black line. Re (B) and *ccdc39* (C) apices 30 days after watering show properly developed sporophytes on Re apices and bundles of gametangia on *ccdc39* apices. D: Phase contrast image of fixed mature *ccdc39* spermatozoids displaying posterior flagellar loops (arrows). E: SEM of *ccdc39* spermatozoid that remains in coils. The nucleus (n) is a long coiled cylinder that narrows posteriorly. Irregular flagella coil around the cell (arrowhead). F: TEM of whole mature spermatozoid in cross section with compacted nucleus (n) in upper coil. Arrowheads indicate irregular flagella that encircle the cell for over 2 revolutions. Cytoplasmic connections (cc) adjoin axonemes across their length. Bar= 500 nm. G-I: TEM cross sections of irregular flagella showing intact acentric central pair complexes (white arrowheads), outer dynein arms (od, black arrows), and double axonemes in single flagellar shafts in G and I.

## Conclusions

Our analyses suggest that the long term vegetative propagation of the Gd lab strain led to accumulation of several (epi-)mutations, affecting the male reproductive apparatus. Unfortunately, the original Gd collection site is not available anymore due to changes in land use. However, the Gd isolate was shipped to Japan (GdJp) in the early 90s and has been propagated involving regular sexual reproduction. To proof the assumption that the original Gd isolate had not been infertile, we analyzed the fertility of GdJp and found that the sporophyte development was significantly higher in comparison with the predominantly vegetatively propagated Gd (Fig. S10). Hence, the genetic and epigenetic differences observed between Gd and Re are probably due to somatic changes acquired during vegetative culture as previously proposed (Ashton *et al.* 2000). To avoid such accumulation of deleterious (epi-)mutations it appears advisable to regularly include sexual reproduction into the lab strain propagation, and to test the offspring for fertility.

Interestingly, known key players affecting cilia motility in mammal model organisms and human could be identified via their *P. patens* expression profile as well as through (epi-)mutations between Gd and Re. Some of these were characterized via transcript profiling, demonstrating biased expression between the ecotypes. Loss-of-function analysis of ccdc39 showed involvement in proper flagella formation in *P. patens* as well. Hence, flagellated plants (like mosses) apparently harbor male gametes with a flagellar architecture that is conserved with mammals, and thus could serve as easily accessible models for human disease or animal foodstock fertility associated with flagella. Whether only flagella or the process of male motile gamete development is evolutionarily conserved as well remains to be determined.

Our network analyses show that the components of the protein-protein interaction network of the eukaryotic flagellar structure are well conserved. This network is also supported by the loss-of-function mutant: ccdc39 was found via the network and expression analysis as well as through a candidate gene approach, and resembles the Gd phenotype in terms of male infertility. Most probably, given the lack of SNPs and DMPs in the underlying gene, Ppccdc39 is not the causative master mutation. Rather, it appears probable that one or several upstream regulators are affected, causing (among others) the mis-regulation of Ppccdc39. In this regard it is interesting to note that the homeodomain (HD) TALE TF BELL1, known to be involved in the alternation of generations (haploid to diploid phase transition) in algae and plants (Horst *et al.* 2016, Lee *et al.* 2008), shows expression in Re but not in Gd. Intriguingly, the heterodimeric interaction of HD TALE TFs observed in animals and plants has recently been suggested to be an ancestral eukaryotic feature to control the haploid-to-diploid transition only when gametes were correctly fused (Joo *et al.* 2018). Potentially, master regulators like BELL1 not only control zygote formation, but even earlier processes of sexual reproduction.

Primary Ciliary Dyskinesia (PCD) is a heterogeneous genetic defect that results in abnormal structure and function of cilia (Noone *et al.* 2004). In humans, the condition has a range of biological and clinical phenotypes that include respiratory infections, reduced fertility and situs inversus or developmental left-right inversion of organs. Ultrastructural observations of cilia in patients with PCD reveal perturbations of outer and inner dynein arms, radial spokes and the central pair complex, with 10% exhibiting normal axonemal structure (Papon *et al.* 2010). The conserved nature of the 9+2 core structure of cilia and flagella is particularly underlined in the present study as a universal phenotype of ccdc39 loss-of-function mutants, since it occurs in moss, dogs and humans (Merveille *et al.* 2011). All show axonemal disorganization and inner dynein arm aberrations. Functional analyses indicate that the ccdc39 gene is necessary to assemble the dynein regulatory complex and inner dynein arms. The striking defects evident in *ccdc39* moss spermatozoids are even more exaggerated than those in humans and dogs, perhaps due to the extreme length of flagellar axonemes compared with cilia. Nevers (Nevers *et al.* 2017) classified motility-associated genes responsible for ciliopathies such as PCD, including ccdc39, in a relatively small subset of ciliary genes necessary to construct a motile flagellum. These genes are found in all ciliated organisms, including eukaryotes such as land plants in which cilia are restricted to gamete-producing cell lineages. As an easily manipulated experimental system, *P. patens* provides unparalleled opportunities to systematically and individually examine gene structure, function and evolution in the core genes responsible for the development of the eukaryotic cilium/flagellum.

## Methods

### Plant material

*P. patens* ecotypes Gd (Rensing *et al.* 2008) and Reute (Hiss *et al.* 2017) were cultivated under the same conditions as described in (Hiss *et al.* 2017) for sporophyte development, and for crossing analysis as described in (Perroud *et al.* 2019). For gametophytic stage phenotyping, protonemal cells were placed on 9cm Petri dishes with solid minimal medium (1% w/v agar, Knop’s medium, (Knop 1868)) enclosed with 3M micropore tape and inoculated 7 days for protonema and in total 10 days for gametophore analysis in long-day (LD) conditions (70 μmol m^−2^ s^−1^, 16h light, 8h dark, 22°C). Gametangia harvest and analysis was performed at 21 days after short-day (SD, 20 µmol m^−2^ s^−1^, 8h light, 16h dark, 15°C) transfer.

### Counting and statistical analysis of sporophytes per gametophore under selfing and crossing conditions, data visualization

To determine the number of sporophytes developed by ecotypes and knock-out mutant, counting of the selfed F1 of three independent mutant lines (*ccdc39#8, ccdc39#41, ccdc39#115*, Fig. S11) was performed at 30 days after watering as described in (Hiss *et al.* 2017) using a Leica S8Apo binocular (Leica, Wetzlar, Germany). Counting of the crosses was performed according to (Perroud *et al.* 2019) using a fluorescence stereomicroscope SteREO LumarV12 (Carl Zeiss, Oberkochen, Germany). For counting of the selfed F1, at least five independent biological replicates with in total 537 to 742 gametophores per sample were analysed. For the crossings, three biological replicates with in total 300 to 630 gametophores per sample were analysed. For the ecotype comparison between Re, Gd and GdJp three replicates with in total 523 to 730 gametophores were analysed. Statistical analysis was carried out using Microsoft Excel 2016 (Microsoft) and R. Plots in R were done using ggplot2 (Wickham 2016).

### Microscopic analysis of gametangia and sporophytes

Harvest and preparation of sporophytes, gametangia and spermatozoids was performed with a Leica S8Apo binocular (Leica, Wetzlar, Germany). Microscopic images were taken with an upright DM6000 equipped with a DFC295 camera (Leica). For both devices the Leica Application suite version 4.4 was used as the performing software Images were processed (brightness and contrast adjustment) using Microsoft PowerPoint.

### Spermatozoid microscopy, DAPI staining and counting

Preparation for 4’,6-diamidino-2-phenylindole (DAPI) staining/flagellar analysis and counting of spermatozoids per antheridium was performed after 21 days of SD inoculation. Per sample, a single antheridium was harvested in 4 µl sterile tap water applied to an objective slide. The spermatozoids were released using two ultra-fine forceps (Dumont, Germany) and the sample was dried at room temperature (RT). Samples were fixed with 3:1 ethanol/acetic acid and after drying spermatozoid nuclei were stained applying 0,7 ng/µl DAPI (Roth, Germany) in tap water. The samples were sealed with nail polish (www.nivea.de). Microscopic images were taken with an upright DM6000 equipped with a DFC295 camera (Leica). Brightness and contrast of microscopy images was adjusted using Microsoft PowerPoint.

### Electron microscopy

For SEM images, mature spermatozoids (21 d after SD transfer) were harvested on an objective slide covered with polilysine. Chemical fixation was performed with 2.5% glutaraldehyde and an ascending alcohol preparation using 30, 50, 70, 90, and two times 100 % of EtOH was performed. Subsequently, the samples were critical point dryed (Tousimis, Rockville, USA) and coated with 20nm gold using a Leica Sputter (Leica, Wetzlar, Germany). The samples were then analysed with a SEM (Philips XL30, Hamburg, Germany). For TEM, adult apices (21 d after SD transfer) were used. Sample preparation was done as previously described with slightly modified infiltration (25% Epon (Fluka^®^ Analytical, USA) in propylene oxide for 2h; 50% Epon in propylene oxide overnight; next day pure Epon and polymerization).

### Nucleic acid Isolation

Genomic DNA for genotyping was isolated with a fast extraction protocol, using one to two gametophores as described in (Cove *et al.* 2009). To determine candidate gene expression levels in juvenile (5 weeks LD growth) and adult (21 days after SD transfer) apices, each at least 25 apices were harvested in 20µl RNA later (Qiagen, Hilden, Germany). Tissue was stored at -20°C, until RNA was extracted using the RNeasy plant mini Kit (Qiagen). For RNA-seq RNA was isolated from 40 antheridia bundles, comprising 4-6 antheridia each, harvested at 21 day after SD transfer and directly stored in 20µl RNAlater. RNA was extracted with a combination of RNeasy plant mini and RNeasy micro kit (Qiagen). The RNeasy plant mini kit was used until step 4. Flowthrough of the QIAshredder spin column was diluted in 0,5 volume 100% EtOH and then transferred to the RNEasy spin columns of the micro kit for further treatment with the micro kit. RNA concentration and quality was analysed with Agilent RNA 6000 Nano Kit at a Bioanalyzer 2100 (Agilent Technologies).

### Real-time PCR (RT-PCR) & quantitative real-time PCR (qPCR)

To determine gene expression in juvenile vs. adult, respectively Gd vs. Re adult apices, RNA was extracted as described above. cDNA synthesis was performed using the Superscript III (SSIII) kit (ThermoFisher Scientific) according to the manufacturers’ protocol but using ½ of the suggested SSIII amount. Primers were designed manually with an annealing temperature +/-60°C and a product length of ca. 300 bp. Single genomic locus binding properties were tested by using BLAST against the V3.3 genome of *P. patens*. Real-time PCR was performed using 5 ng of cDNA as input for PCR reaction with OneTaq from New England Biolabs). PCR products were visualized via gel electrophoresis using peqGREEN from (VWR, Germany). As size standard, the 100 bp ladder (NEB) was used. Real-time qPCR was carried out with two (for juvenile vs. adult comparison) or three (expression determination of ccdc39) biological replicates as published in Hiss et al., 2017. As reference gene, act5 (Pp3c10_17070V3.1, (Le Bail *et al.* 2013) was chosen, due to homogenous expression in juvenile and adult apices in Gd and Re (Fig. S12). For primer sequences see Table S10.

### Antheridia bundle RNA-seq & analysis

RNA-library preparation and RNA-seq was performed at the MPI genome center Cologne (mpgc.mpipz.mpg.de). For each ecotype, two libraries were prepared with the ultra low input RNA-seq protocol followed by sequencing with Illumina HiSeq3000 (150 nt, single ended). For gene expression and DEG analysis for Gd 53,6 Mio. and for Re 56,4 Mio. uniquely mapped reads were used. The analysis was performed as previously described for the *Physcomitrella patens* gene atlas project (Perroud *et al.* 2018), using the strict consensus of three DEG callers, defining transcripts as expressed if their RPKM was greater than or equal to 2. For RPKM data see Table S11 and for DEGs Table S8.

### Methylated and differentially methylated positions (DMP) and overlap with single nucleotide polymorphisms (SNP)

The Re data was generated and treated as described for Gd in (Lang *et al.* 2018) and only positions with a coverage equal to 9 or above were used for further analysis. DMPs between Gd and Re were called using methylKit using a q-value < 0.01 and a methylation difference of at least 25% (Akalin *et al.* 2012). Differentially methylated positions were associated with gene models by using bedtools intersect (Quinlan *et al.* 2010). For GO bias calculation all hyper- and hypo-methylated genes within all contexts (CHG, CHH, CG) were used. This gene list was used to identify genes also affected by a SNP between Gd and Re (Hiss *et al.* 2017) to generate GO bias data of the intersection.

### Vector construction and preparation

Deletion constructs were built using pBHRF (Schaefer *et al.* 2010) as backbone. PCR reactions were performed with OneTaq (NEB), restriction enzymes were purchased from NEB as well as the T4 ligase and 100bp respectively 1kb ladder. For primer sequences see Table S1. 5’- and 3’-homologous regions of Ppccdc39 were amplified from gDNA using primer pairs ccdc39_HR1_for/rev and ccdc39HR2_for/rev. Flanks were subsequently cloned into pMETA via TA cloning to generate 5’- and 3’ flank containing vectors. The 5’ flank was cut and cloned into pBHRF using the restriction enzymes *Pme*I and *Xho*I. The 3’ flank was cut and cloned using the restriction enzymes *Nar*I and *Asc*I. The final deletion cassette consists out of the two homologous regions flanking the 35S:Hygromycin:CamVter selection marker (Fig. S13). Plasmid amplification was performed in *E. coli* Top10 cells, plasmid extraction was performed using the NucleoBond Xtra midi kit (Machery-Nagel, Düren, Germany). For stable transfection, the plasmid was cut using *Pme*I and *Asc*I, and after ethanol-precipitation solved in sterile TE.

### Moss transfection

Moss transfection was carried out in the Re wild type background (Hiss *et al.* 2017) with small modifications as published in (Cove *et al.* 2009): after protoplast regeneration all subsequently used media was Knop medium supplemented with 5mM di-ammonium tartrate (Sigma-Aldrich, Germany). For selection, the antibiotic hygromycin B (Roth, Germany) was used as an additive, with a concentration of 20mg/l.

### Genotyping

To identify correctly integrated mutants a set of PCR controls was performed (Fig. S14). WT locus absence was tested by using the primer pair ccdc39_ingDNA_for/rev located in the coding sequence of ccdc39. Proper insertion of the 5’ flank was tested using ccdc39_HR1in (located upstream of the 5’ flank) and p35S_rev (located in the resistance cassette promotor). The 3’ flank was tested using ccdc39_HR2in (located downstream of the 3’ flank) and tCMV_for (located in the terminator of the resistance cassette). Full length PCR was performed using ccdc39_HR1in and ccdc39_HR2in.

### Gene Ontology analysis and visualization

Gene ontology analysis was performed as described in (Widiez *et al.* 2014) based on the *P. patens* V3.3 annotation. Visualization was done using Wordle (http://www.wordle.net/). Green font colors mark over-represented, red font colors mark under-represented GO terms. Size and color correlate with significance. Darker colors and larger font size are correlated with higher significance (q < 0.0001) whereas lighter colors and smaller font size indicate a lower *q*-value (q > 0.0001 but < 0.05).

### Network analysis and gene identifier conversion

For network analysis www.string-db.org (Szklarczyk *et al.* 2015) was used, using V3.3 protein models. To convert gene IDs from the most recent V3.3 annotation to V1.6 used by string-db.org, the conversion table published by (Perroud *et al.* 2018) was used.

## Authors’ contributions

RM and SAR wrote the manuscript with contributions by KSR. RM carried out the experiments and analysed the data. SAR conceived of and supervised the project. PFP helped with the crossing experiments. TH supervised and did the sample preparation for TEM analyses, KSR performed TEM and analysed the images together with RM. FH and RM analysed the RNA-seq data, LS and RM the bs-seq data.

## Acknowledgements

We thank Eva Bieler of the Swiss Nanoscience Institute (SNI), Nano Imaging, University of Basel, for the help taking the SEM pictures, as well as Marion Schön for TEM sample preparation and Rebecca Hinrichs and Jason Backes for assistance. We also thank Karl Ferdinand Lechtreck (University of Georgia) for his helpful comments with regard to the hydin phenotype. The authors declare no conflicts of interest.

## Data availability

All sequencing data has been uploaded to the NCBI SRA. Bs-seq Gd data: SRR4454535 (Lang *et al.* 2018); bs-seq Re data: SRR9901085. RNA-seq data BioProject PRJNA559055.

