## supplemental_figures for "Characterization of evolutionarily conserved key players affecting eukaryotic flagellar motility and fertility using a moss model"

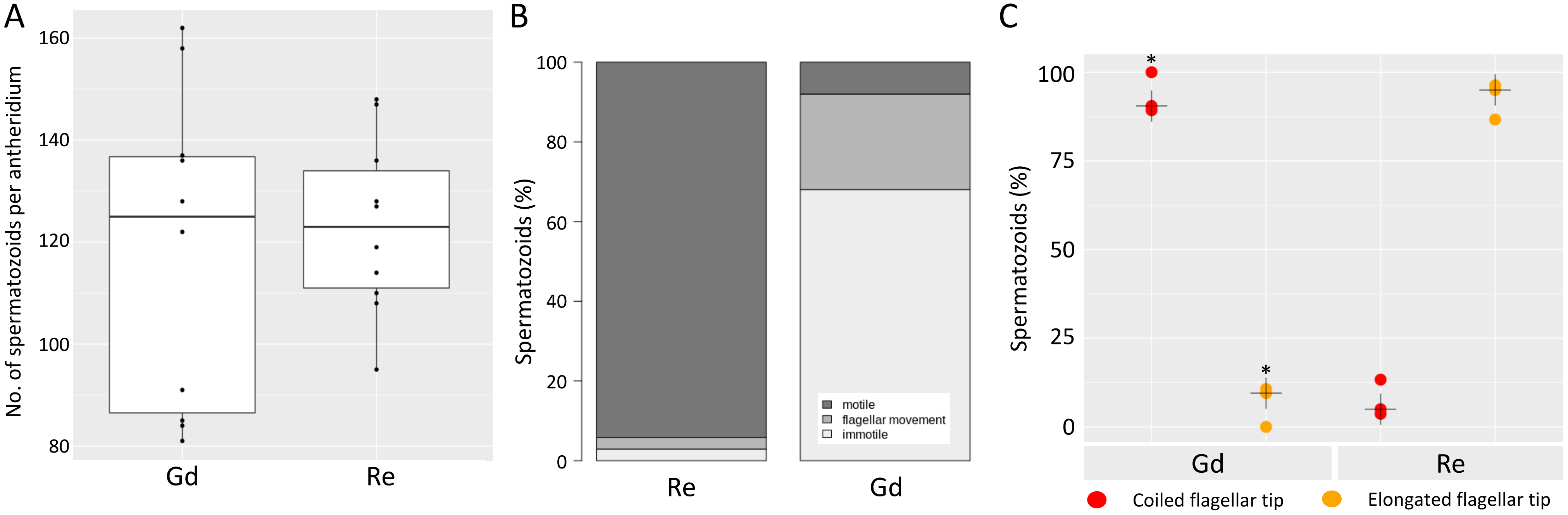


**Figure S1: Detailed analysis of Gd and Re spermatozoids.**

The number of spermatozoids per antheridium does not vary significantly between Gd and Re (A, n = 10 of each ecotype). Median-centered box-dotplots representing 50% of the measurements within the white box, whereas the whiskers show the 1.5 interquartile range (IQR). Dots show individual measurements. In Re, 94% of the spermatozoids were motile, 6% were immotile of which 3% showed flagellar movement. In Gd, 8% of the spermatozoids where motile, 92% were immotile, of which 24% showed flagellar movement. The number of motile spermatozoids is significantly different between ecotypes (B, Gd n = 50, Re n = 103, $p < 0.01$ chi-square test). C: Gd spermatozoids (n = 51) show coiled flagellar tips statistically significantly more often compared to Re spermatozoids (n = 62, $p < 0.01$, t-test). Median: black cross.

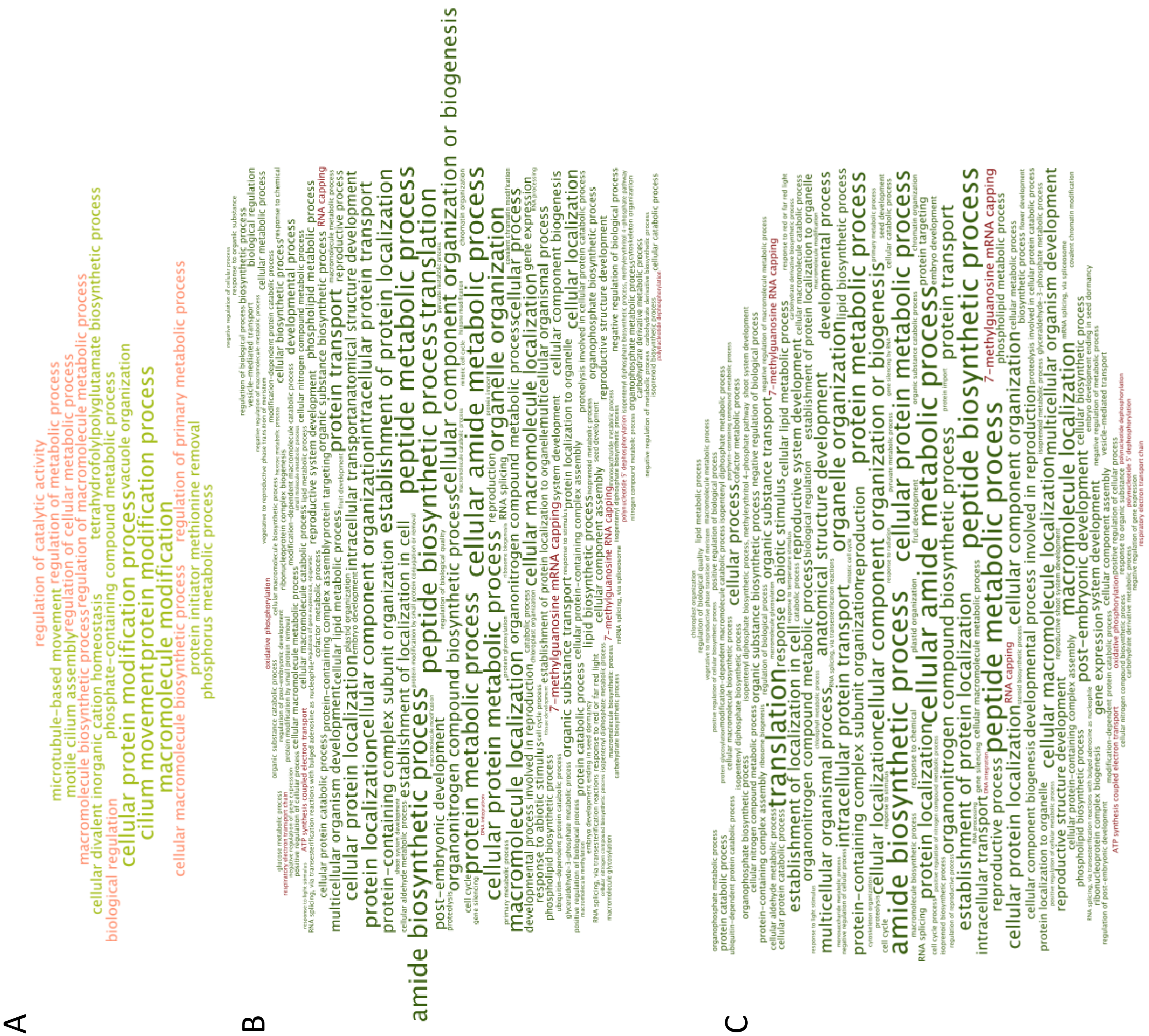


**Figure S2: GO bias analysis and word cloud vizualization.** Over-represented terms are shown in green, whereas under-represented terms are shown in red. Larger font size correlates with a higher significance level. Genes affected by DMPs between Re and Gd in any of the contexts (CG, CHH, CHG) show over-representation of terms connected to cilia motility as well as terms related to macromolecule modification (A). Genes expressed in Gd (B) and Re (C) antheridia bundles show over-represenetation of polyamine and peptide biosynthetic processes as well as under-represented terms associated with mRNA capping.

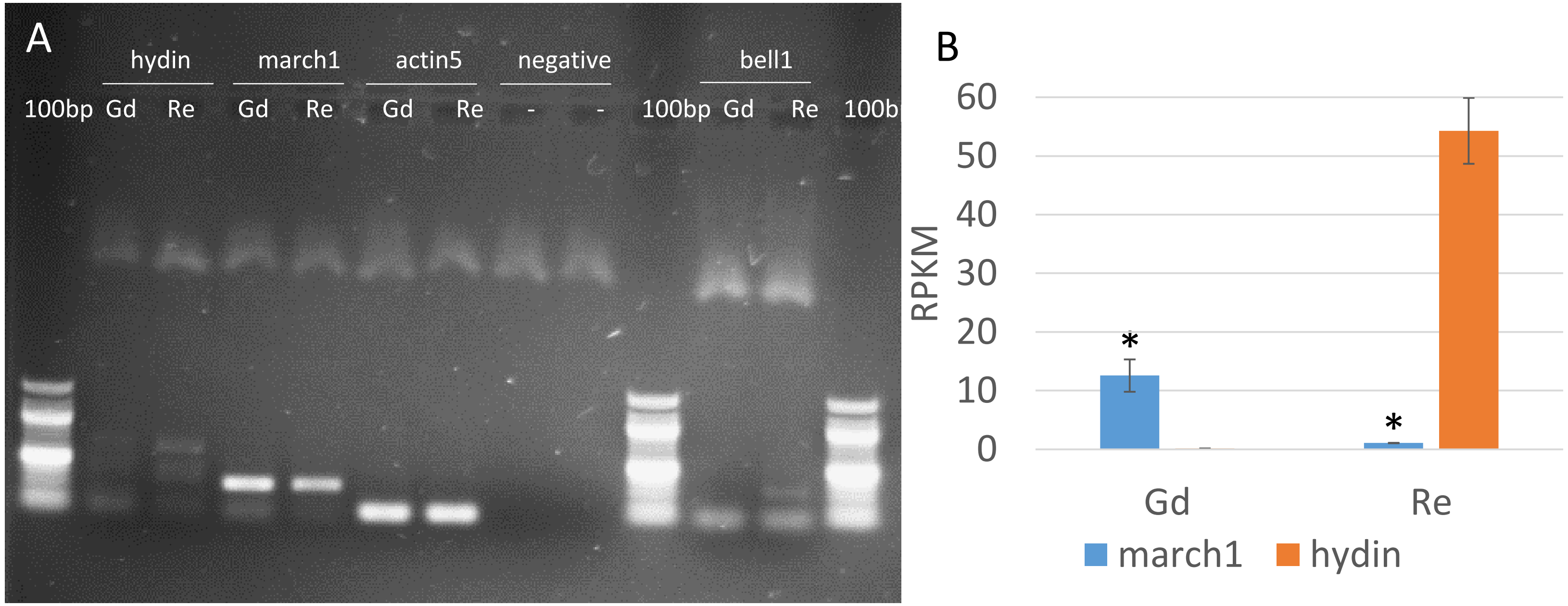


**Figure S3: Expression analysis of genes expressed in antheridia bundles.**

A: Expression level in adult gametophore apices of Gd and Re of hydin, march1 and bell1 genes were verified via RT-PCR. B: RNA-seq expression of Gd and Re march1 (blue) and hydin (orange) show significant differences between Gd and Re (*, $p < 0.01$ t-test).

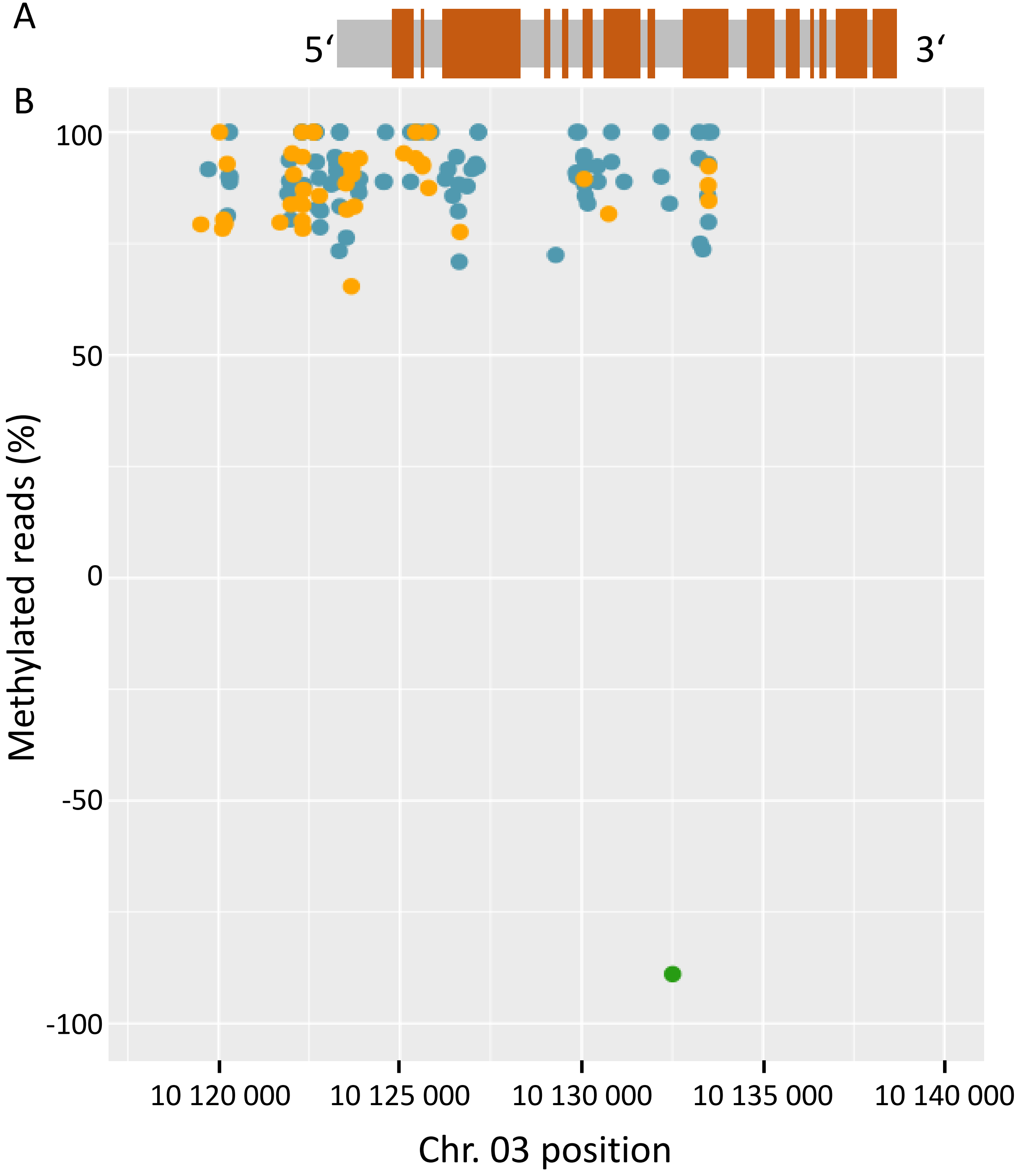


**Figure S4: Hydin gene.**

A: Exon and intron pattern of hydin. B: Aligned with the differentially methylated positions (DMPs) whereas 0 to 100 represents positions methyated with 0-100% in Gd and 0 to -100 represents positions methylated with 0-100% in Re. Gene body and 5‘-UTR of Gd hydin are highly affected by DNA methylation showing CHG (94, blue) and CHH (41, yellow) methylation, whereas in Re a single CG (green) methylation could be detected in an intron.

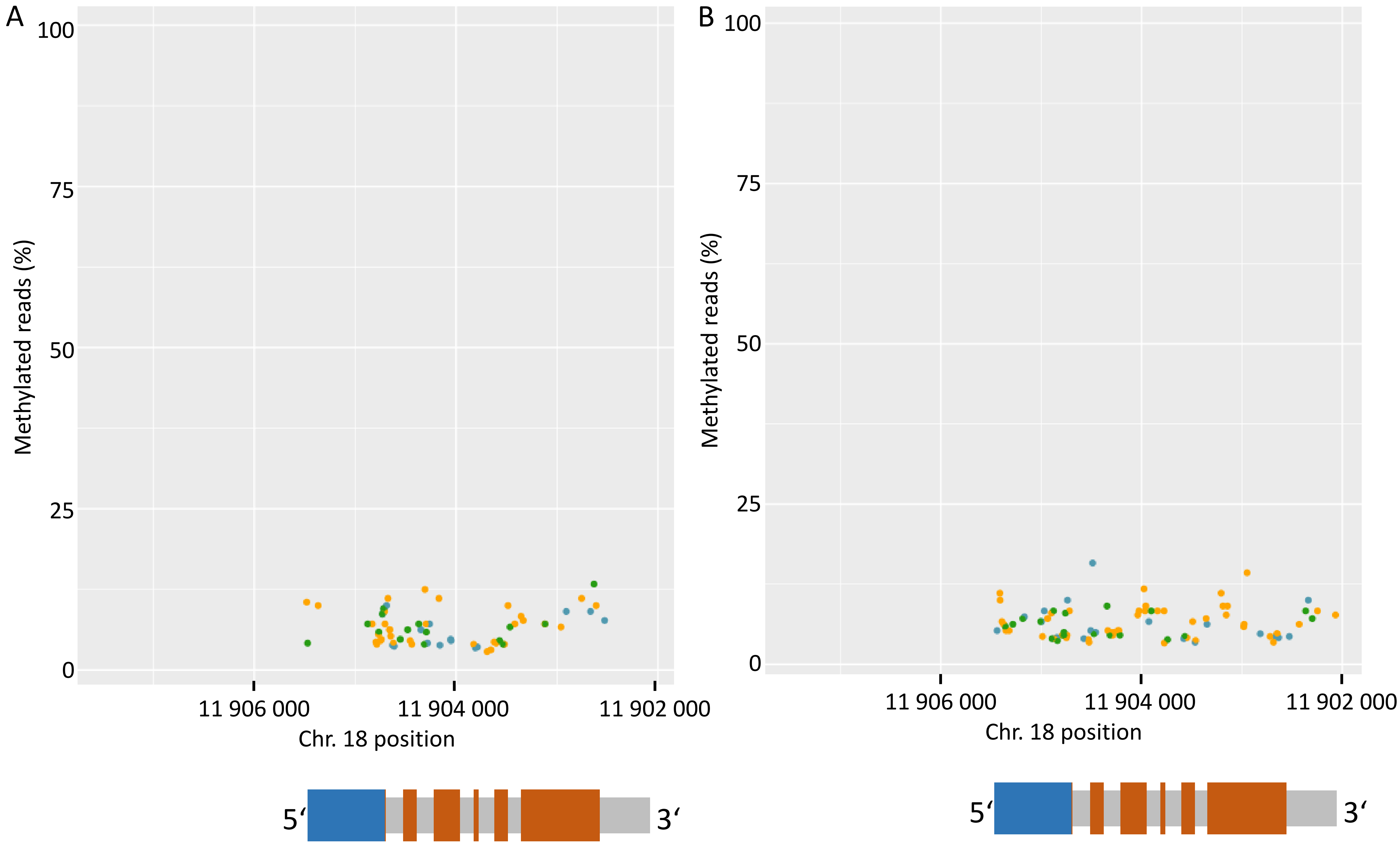


**Figure S5: March1 gene body methylation.**
Percentage of methylated reads per position per pattern of Gd (A) and Re (B) adult gametophores. The V3.3 gene model of march1 is shown in grey, the UTRs are marked in blue and exons are shown in orange. Very low levels of methylation are detectable in march1 and 2kb upstream in both Gd and Re. CHG (blue), CHH (yellow) and CpG (green) marks are shown.

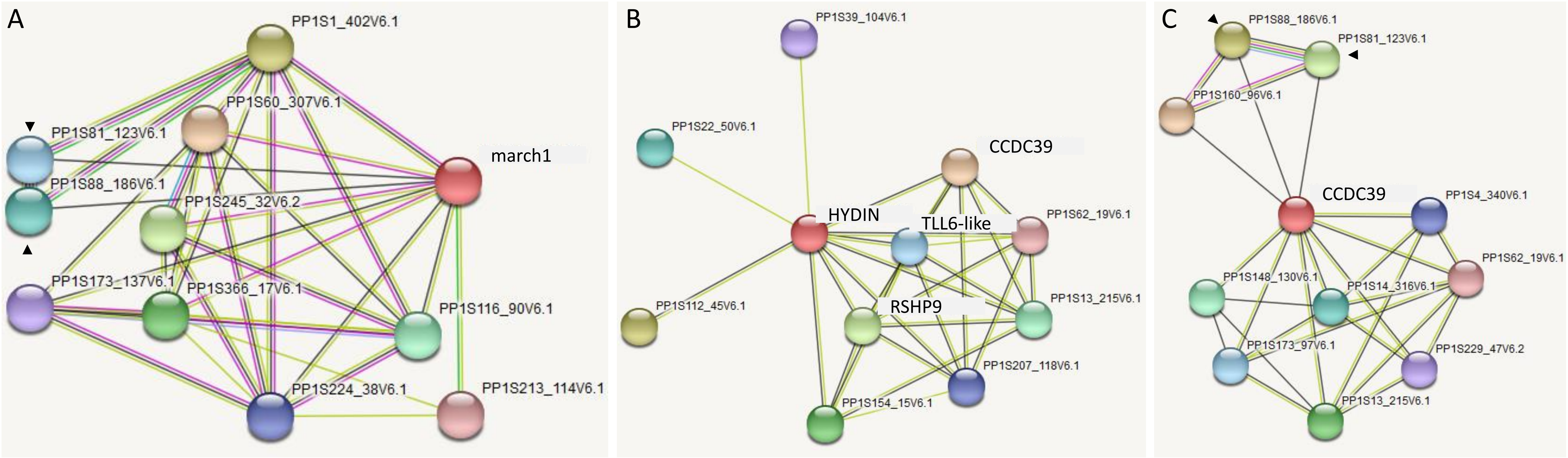


**Figure S6: Network analysis of genes associated with proper flagellar functionality.**

A: March1 network shows two protein phosphatases being co-expressed in common with CCDC39 (C, black arrows). B: Network analysis of hydin shows connectivity with genes affecting sperm cells (CCDC39, RSPH9, TLL6-like). C: Network analysis of CCDC39. Coexpressed protein phosphatases marked by black arrows. Line color specifies connection type between analysed proteins: dark grey marks co-expression, light green marks literature analysis, sky blue marks protein homology, grass green marks gene neighborhood, cyan marks interaction shown by curated databases, pink marks experimentally determined interactions.

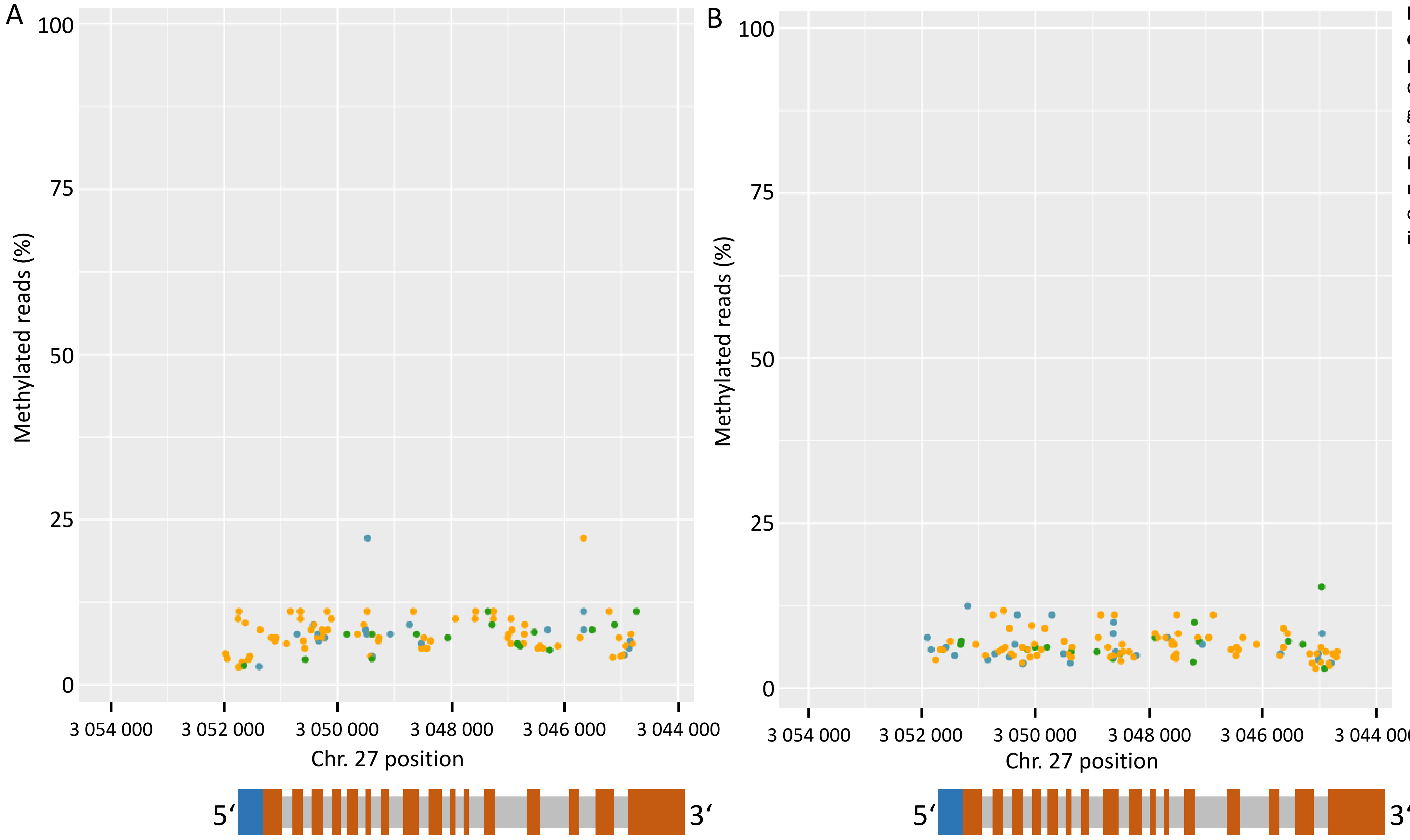


**Figure S7: Percentage of ccdc39 methylated reads per position.**
Gd (A) and Re (B) adult gametophores. Gene model annotation as described in Fig. S5. Low gene body methylation is detectable in ccdc39 and 2kbp upstream in both Gd and Re.

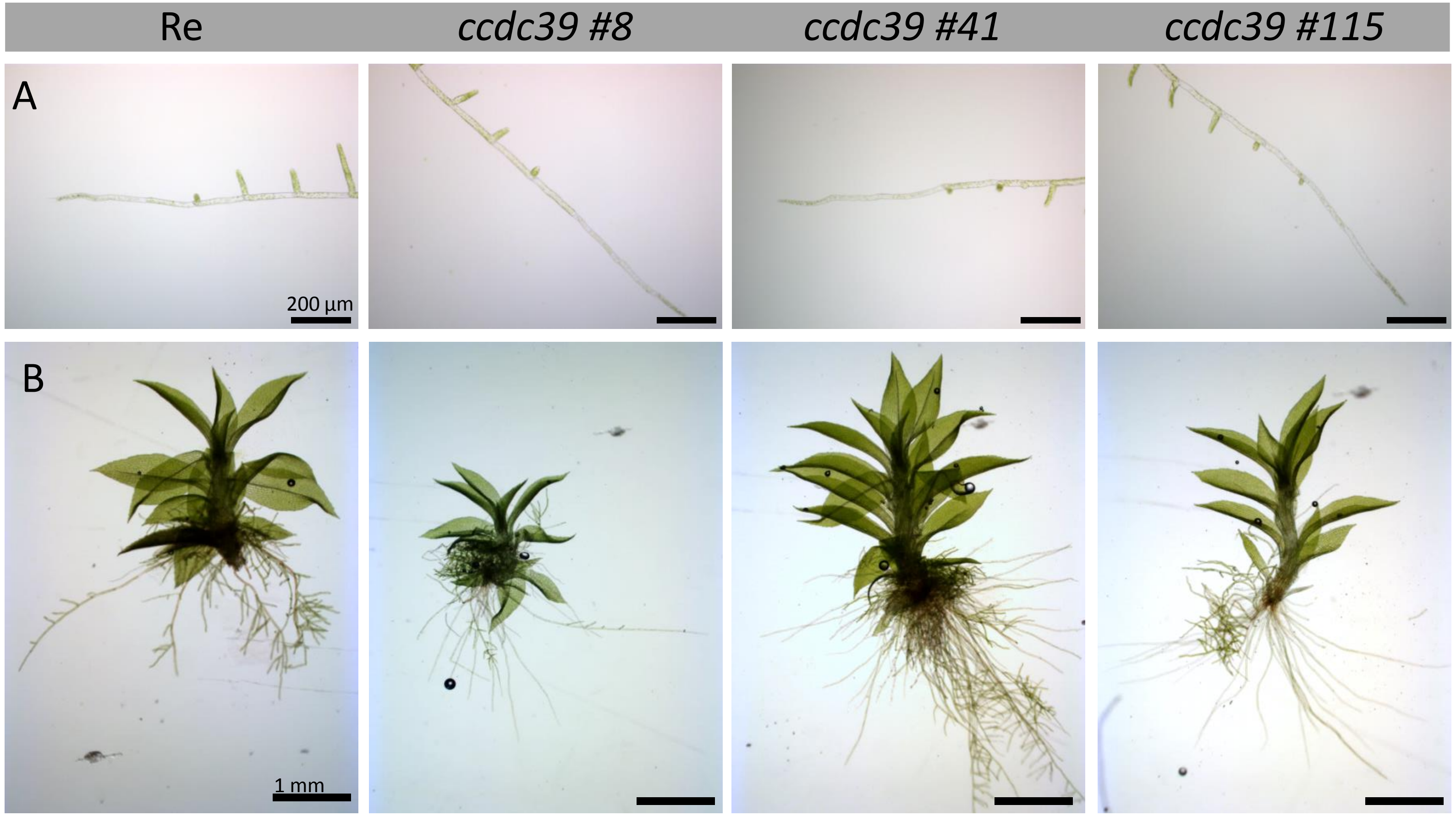


**Figure S8: Protonema (A) and gametophore (B) development of Re compared to *ccdc39* mutant strains**.
No obvious morphological differences could be observed in the analysed mutant lines.

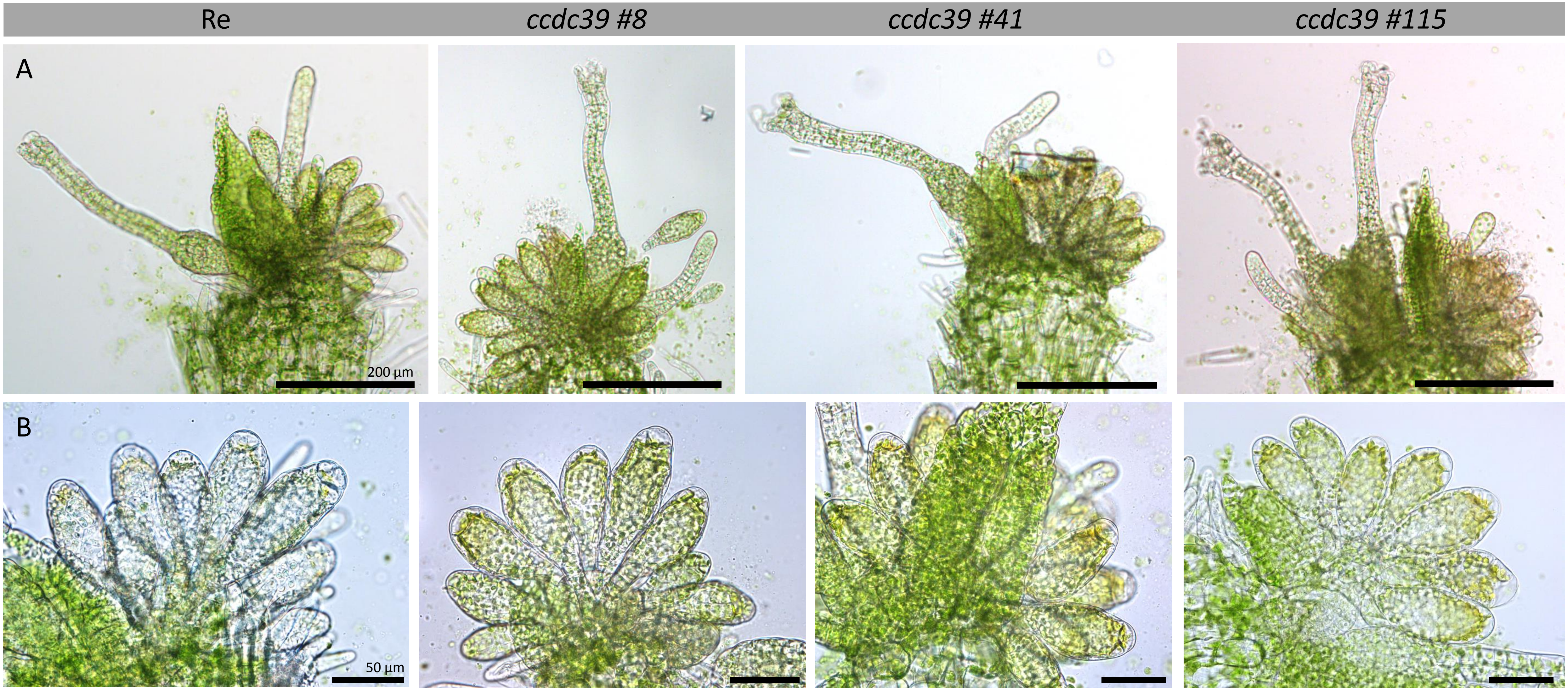


**Figure S9: Gametangia development of *ccdc39* mutant strains compared to Re.**
No obvious morphological differences can be detected. A: dissected apices with both archegonia and antheridia, B: detailed images of the antheridia bundles.

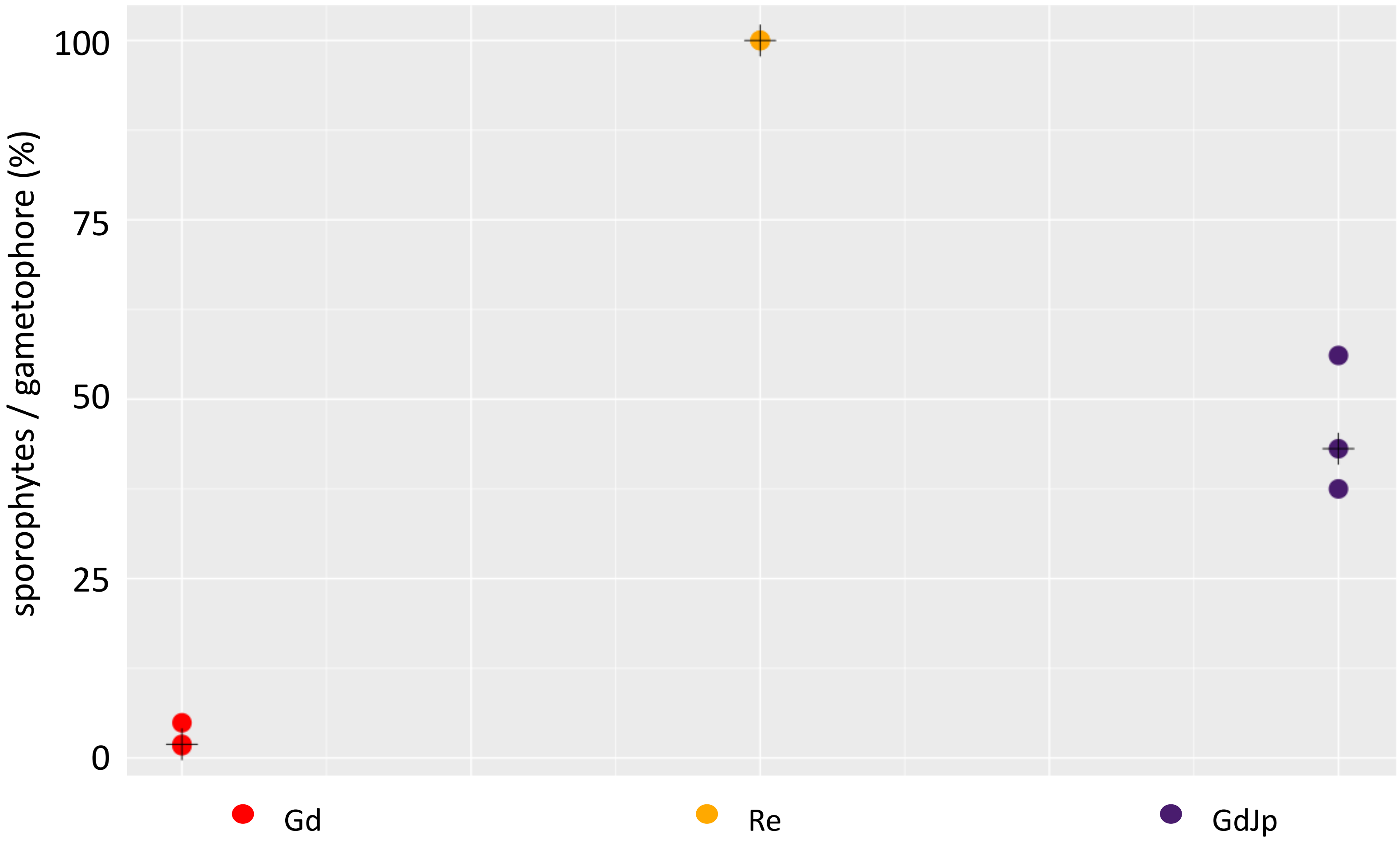


**Figure S10: Number of sporophytes per gametophore developed under selfing conditions for *P. patens* ecotypes Re, Gd and GdJp.**

Re (n = 3, 730 gametophores) shows median 100% sporophyte per gametophore, Gd (n = 3, 523 gametophores) shows 1.9% sporophytes per gametophore and GdJp (n = 3, 653 gametophores) 43.1%. GdJp develops significantly more sporophytes per gametophore in comparison with Gd ($p < 0.05$, t-test).

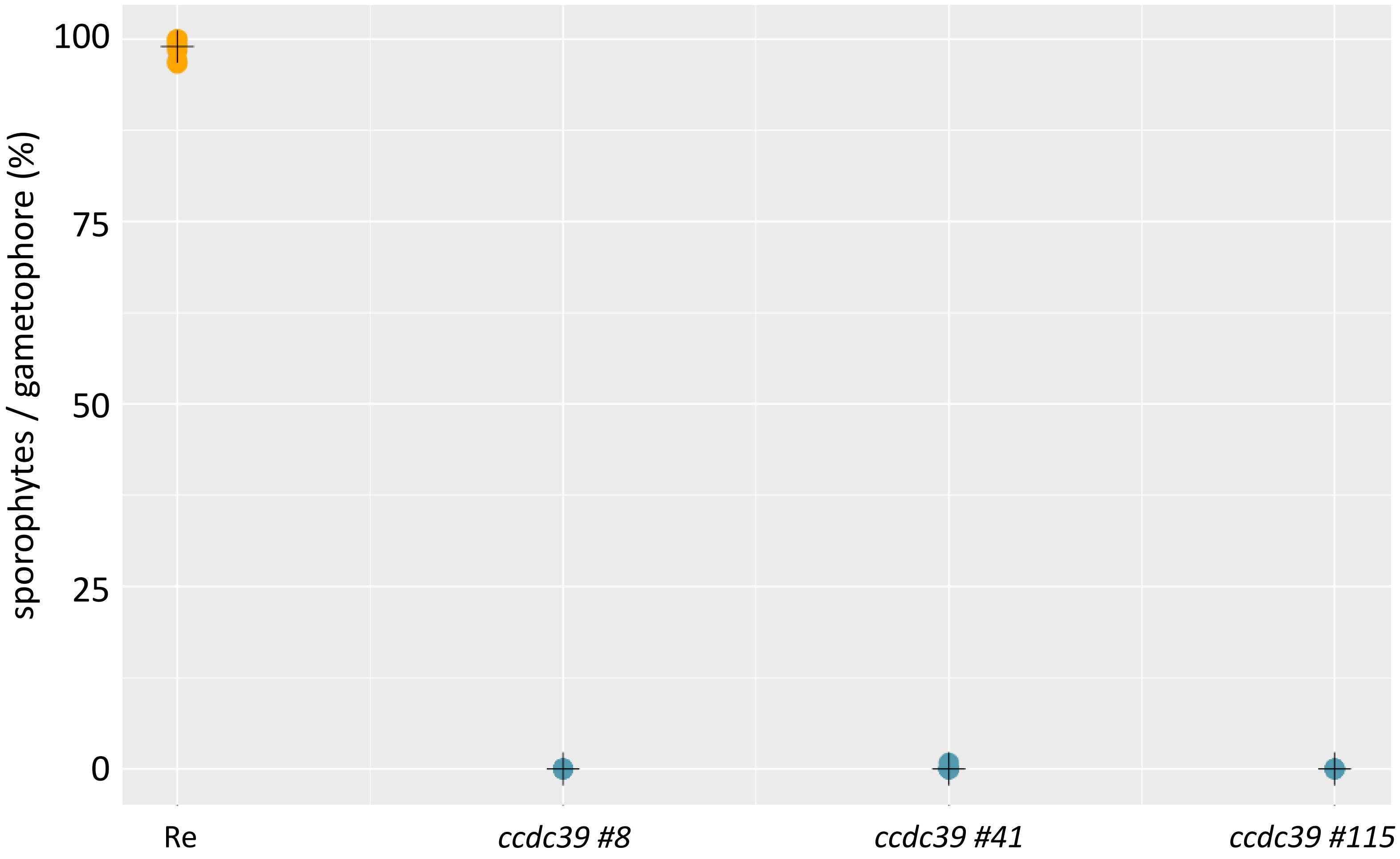


**Figure S11: Sporophyte per gametophore development for all three independently analysed *ccdc39* strains.**
All *Ccdc39#8* (n = 546), *Ccdc39#41* (n = 714) and *Ccdc39#115* (n = 537) show in median 0% of sporophytes in comparison to median 99% of sporophytes for the corrosponding wildtype background Re.

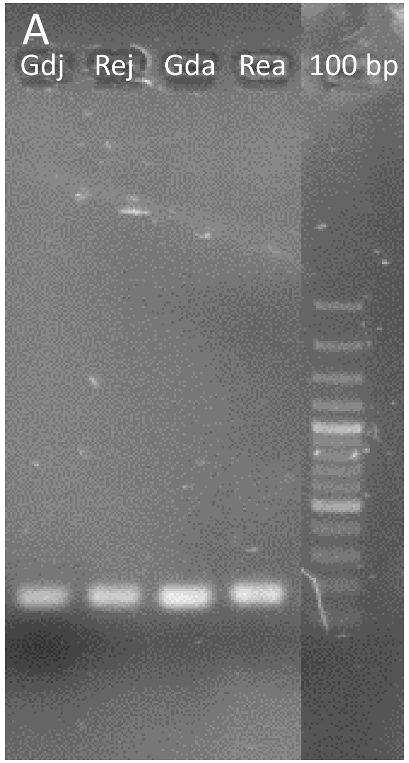


**Figure S12: Expression level of act5 in juvenile (j) and adult (a) apices of Gd and Re show similar expression.**

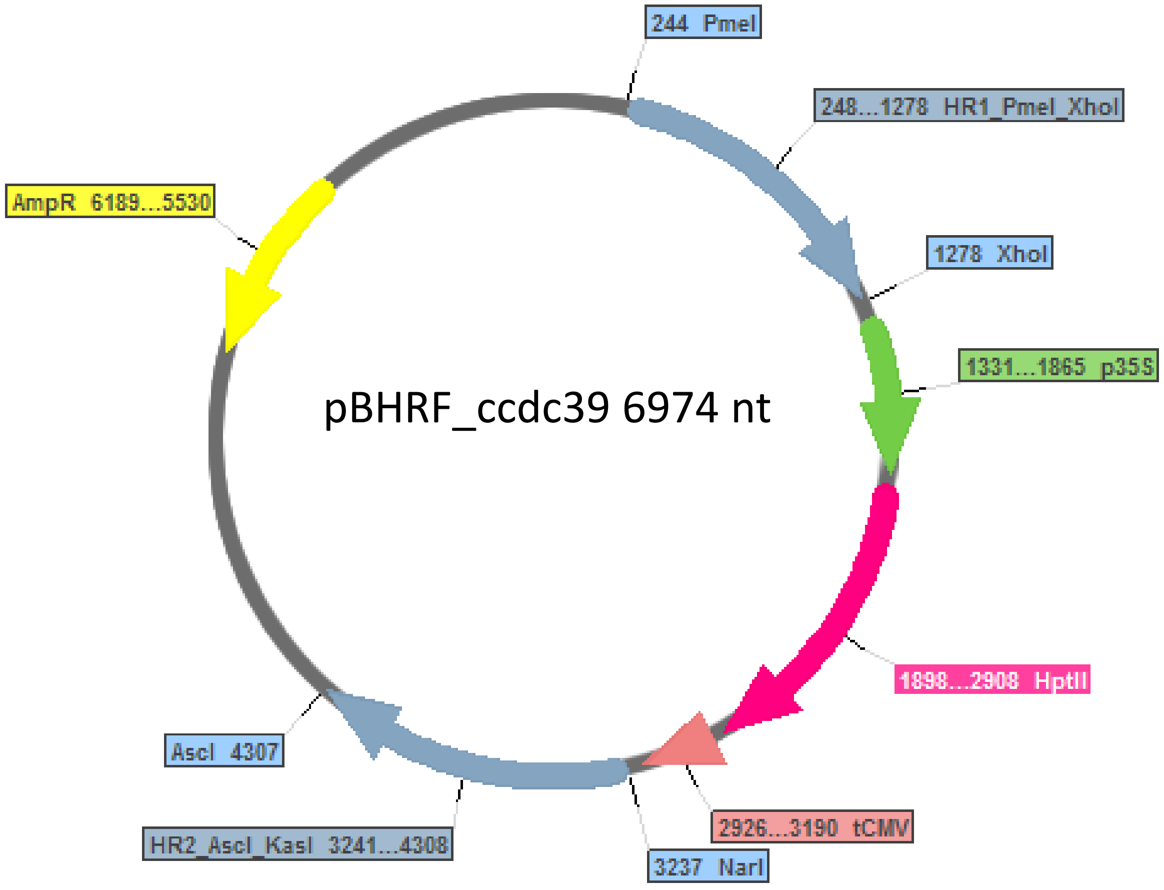


**Figure S13: Final vector for amplification via ampicillin selection (yellow) of the knock out cassette flanked by the enzymes *Pme*I and *Asc*I.**
HR1 and HR2 (blue) flank the resistance cassette consisting out of the 35S promotor (green), the *hptII* resistance gene (pink) and the CMV terminator (red).

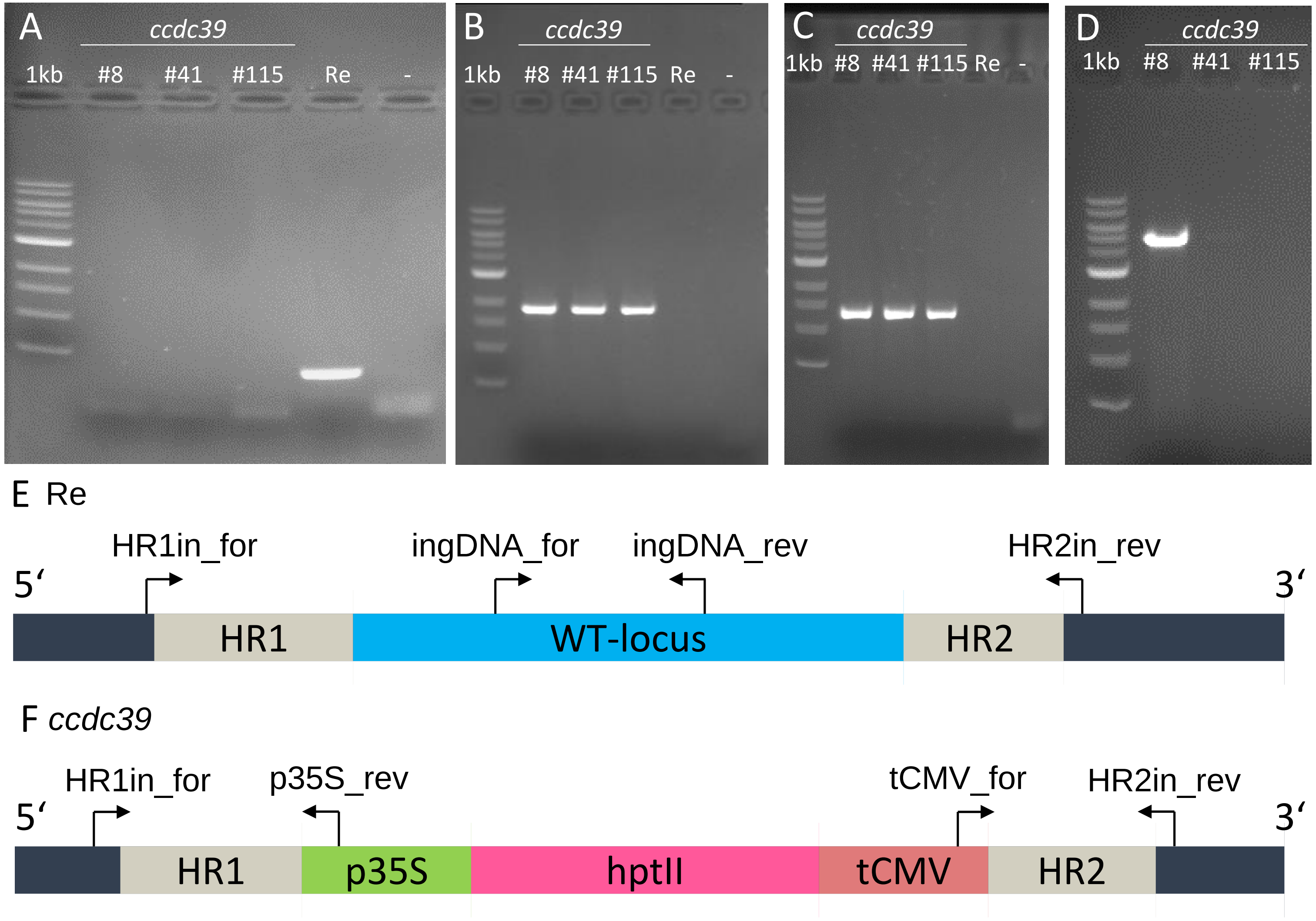


**Figure S14: Gel images and sketch (E,F) of performed genotyping on Re and *ccdc39* gDNA.**
Presence of the wild type locus was tested using ingDNA_for/rev (A). HR1 (B) and HR2 (C) presence was verified using HR1in_for/p35S_rev and tCMV_for/HR2in_rev. Full length amplification was performed using HR1in_for/HR2in_rev (D).
